# Extensive rewiring of the gene regulatory interactions between in vitro-produced conceptuses and endometrium during attachment

**DOI:** 10.1101/2023.08.03.551863

**Authors:** Fernando H. Biase, Sarah E. Moorey, Julie G. Schnuelle, Soren Rodning, Marta Sofia Ortega, Thomas E. Spencer

## Abstract

Pregnancy loss is a significant problem when embryos produced *in vitro* are transferred to a synchronized uterus. Currently, mechanisms that underlie losses of *in vitro-produced* embryos during implantation are largely unknown. We investigated this problem using cattle as a model of conceptus attachment by analyzing transcriptome data of paired extraembryonic membrane and endometrial samples collected on gestation days 18 and 25, which spans the attachment window in cattle. We identified that the transfer of an *in vitro-produced* embryo caused a significant alteration in transcript abundance of hundreds of genes in extraembryonic and endometrial tissues on gestation days 18 and 25, when compared to pregnancies initiated by artificial insemination. Many of the genes with altered transcript abundance are associated with biological processes that are relevant to the establishment of pregnancy. An integrative analysis of transcriptome data from the conceptus and endometrium identified hundreds of putative ligand-receptor pairs. There was a limited variation of ligand-receptor pairs in pregnancies initiated by *in vitro-produced* embryos on gestation day 18, and no alteration was observed on gestation day 25. In parallel, we identified that *in vitro* production of embryos caused an extensive alteration in the co-expression of genes expressed in the extraembryonic membranes and the corresponding endometrium on both gestation days. Both the transcriptional dysregulation that exists in the conceptus or endometrium independently, and the rewiring of gene transcription between the conceptus and endometrium are a potential component of the mechanisms that contribute to pregnancy losses caused by in vitro production of embryos.

**SIGNIFICANCE STATEMENT:** The successful establishment of pregnancies following the transfer of an *in vitro* produced embryo is essential for cattle production and assisted human reproduction. Most of the pregnancies initiated by the transfer of an *in vitro* produced embryo fail, in part because of dysfunctional interaction between the embryo and endometrium during pregnancy establishment. Our study identified that conceptuses produced *in vitro* and their corresponding endometrium have massive dysregulation in gene activity during the peri-implantation window, which affects crucial biological functions necessary for pregnancy. These gene expression alterations are a major contributor to the high rates of pregnancy loss following the transfer of an *in vitro* produced embryo. Our findings have implications for improving assisted reproduction in both agriculture and biomedicine.

## INTRODUCTION

Successful attachment or implantation of the conceptus (embryo and associated extraembryonic membranes) into a receptive uterus is an important milestone for pregnancy in cattle and other mammals. In cattle, by gestation days 4-5, the blastocyst enters the uterine lumen and sheds its zona pellucida, allowing for direct contact between the outer monolayer of trophectoderm cells and the uterine luminal epithelium (LE) (1-3). At this time, the blastocyst begins to produce interferon tau (IFNT) (4), which is the major pregnancy recognition signal that inhibits the development of the endometrial luteolytic mechanism (5, 6). On gestation days 12-14, the blastocyst is ovoid in shape (∼2-5 mm in length) and transitions into a tubular shape by days 14-15, at which time it can be termed a conceptus (7). In cattle, there is a significant release of IFNT as the conceptus begins to elongate around day 15 (8). The conceptus elongates via the proliferation of the trophectoderm and parietal endoderm cells (9) and reaches 20 cm or more in length by day 19-20 (9, 10). The trophectoderm begins to attach to the endometrial lining (9, 11) thereby initiating cell-to-cell communication mediated by adhesion (11). By gestation day 25, a small proportion of trophectoderm cells have differentiated into binuclear cells (12), and the formation of the chorion marks the onset of the epitheliochorial placentation (10, 13, 14).

Alongside INFT, blastocysts produce a series of bioactive molecules named embryotropins (15) that can have autocrine or paracrine effects. Paracrine effects are observed in the endometrium as early as gestation day 7 when the embryo modulates the regulation of gene expression in endometrial areas surrounding it (16), also causing alterations in the metabolite composition of the uterine luminal fluid (17). Global alterations of the gene expression in the endometrium are observed during elongation by gestation days 15-16 (18-20). As the trophoblast and endometrium lining have direct contact, cell-to-cell interactions can be established through ligand-receptor mediated signaling (20-22), and ultimately there is a synchrony of gene regulation between the conceptus and endometrium (23).

In cattle, most conceptus losses (70%-80%) occur before pregnancy is recognized and placentation starts (24-29). These losses can be accentuated when an *in vitro* produced embryo is transferred to a surrogate dam (30-34). *In vitro* produced conceptuses present altered gene expression during the attachment phase (35-37), and this dysregulated gene expression influences the regulation of genes in the endometrium (22, 38-40). The molecular interaction between conceptus and endometrium is an important feature during attachment (23), and dysregulation of conceptus-maternal communication is a probable cause for lower pregnancy establishment rates following the transfer of *in vitro* produced embryos. Therefore, *in vitro* produced embryos are a model to understand the dysregulated biological pathways associated with pregnancy interruptions at implantation or pathological placentation.

In the current study, we aimed to determine whether there are differences in transcript abundance in extraembryonic membranes and the harboring endometrium in the peri-attachment period in pregnancies initiated by the transfer of an *in vitro*-produced embryo or artificial insemination. Overall, we hypothesized that conceptuses produced *in vitro* disrupted gene regulatory interactions with the harboring endometrium in the peri-attachment period. Our results revealed major alterations in transcript abundance in extraembryonic membranes and endometrium of gestations carrying an *in vitro* produced conceptus. This disrupted regulation of gene expression causes the conceptus and the endometrium to rewire their gene regulatory interactions. The combination of dysregulated gene transcription and the newly created blueprint of interaction between conceptus and endometrium can drive pregnancy loss during peri-attachment in pregnancies initiated by the transfer of an *in vitro*-produced embryo.

## RESULTS

### Experiment overview

We collected samples from extraembryonic and endometrial tissues from pregnancies initiated by artificial insemination or transfer of an *in vitro* produced embryo with gestations interrupted on day 18 or 25. Representative pictures are shown in Fig. 1A. We dissected day 18 conceptuses into extraembryonic tissues (EET) and embryonic disk, but only EETs were used for subsequent analysis. We also dissected day 25 conceptuses into the conceptus proper and EETs. The EETs were further separated into chorion, allantois, or undifferentiated EET; but only samples from chorionic tissue were used for analysis. We collected samples from the caruncular and inter-caruncular areas of the endometrium (∼2mm deep sampling) separately. Altogether, we produced RNA-sequencing data from 14 pregnancies interrupted on gestation day 18 and 13 pregnancies interrupted on gestation day 25 (N _day 18 artificial insemination_: 7, N _day 18 in vitro produced embryos_: 7, N _day 25 artificial insemination_: 7, N _day 25 in vitro produced embryos_: 6, Fig. 1B). Over 3.4 billion reads were mapped to the genome and matched to the Ensembl annotation (SI Appendix, Table S1), which allowed us to estimate the transcript abundance of 12,180 and 14,047 genes (protein coding, long non-coding RNA, and pseudogene) for EET and endometrial samples, respectively (Fig. 1B).

**Fig. 1.**
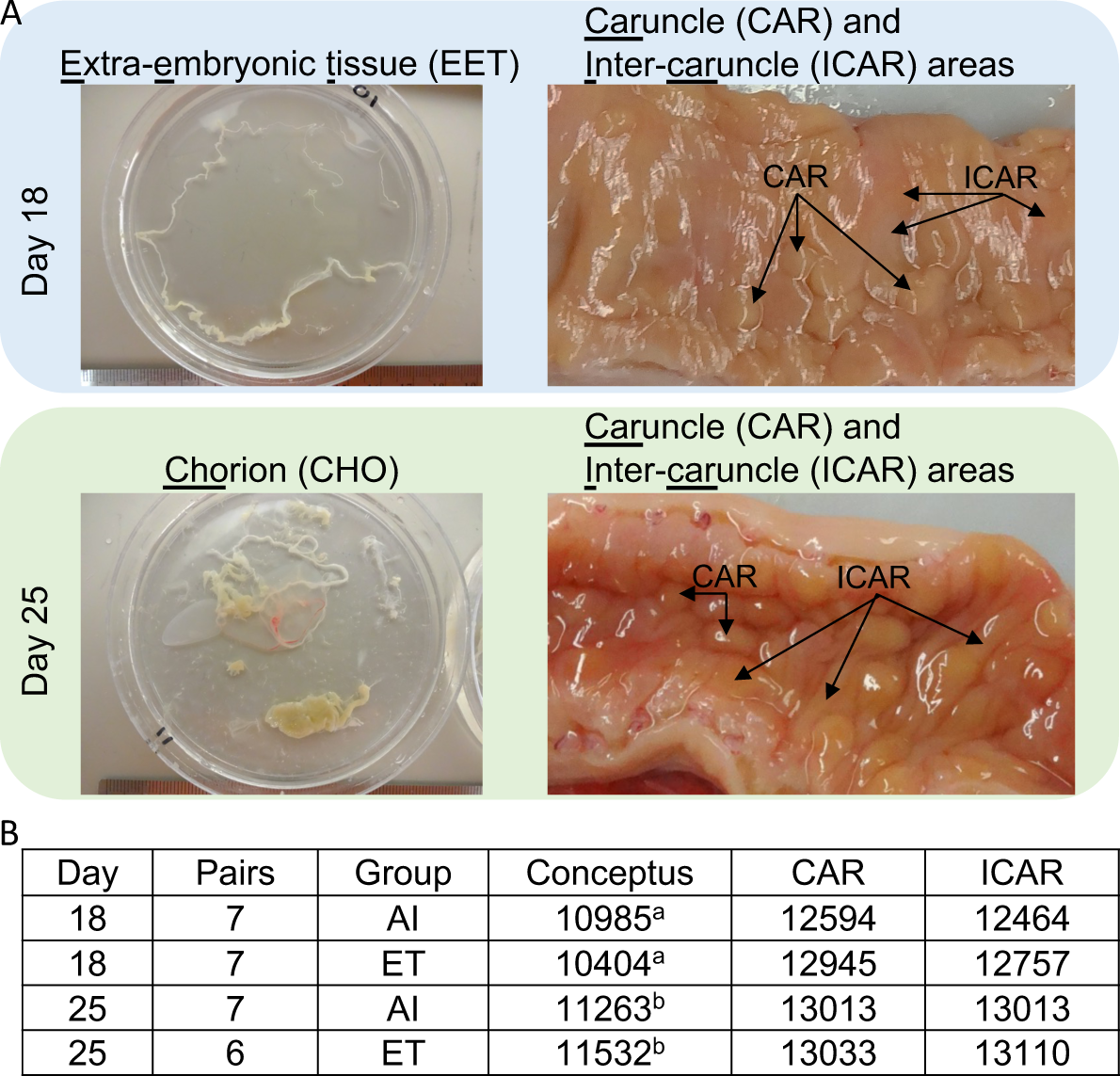
Overview of the biological samples and transcriptome data produced. (A) Representative images of the conceptuses and endometrial tissues that were sampled on gestation days 18 and 25. Uterine sections show the endometrium after sectioning the uterine wall on the vertical axis. (B) Sample size and the number of genes obtained per tissue type. ^a^: extraembryonic tissues, ^b^: chorion. AI: artificial insemination; ET: embryo transfer.

The large number of samples analyzed requires us to present only a subset of the results that are most relevant to the biology of the conceptus-maternal interaction. In the main text, we focused on the differences in the transcriptome related to the gestation progress (day 25 versus day 18 in pregnancies initiated by artificial insemination) and comparisons of the samples collected from pregnancies initiated by either artificial insemination or the transfer of an in vitro-produced embryo. The remainder of the analyses (day 25 versus day 18 in pregnancies initiated the transfer of an in vitro-produced embryo, as well as an interaction between gestation time and groups) can be found in the supplementary text (SI Appendix).

### Changes in the transcriptome of extraembryonic tissues and endometrium between gestation days 18 and 25 in gestations initiated by artificial insemination

We inferred 3376 genes with differential transcript abundance between day 18 EET and day 25 chorion (FDR<0.01, Fig. 2A, Dataset S1). Approximately 43% of these genes were exclusively expressed in either of those days of development (Fig. 2A), whereas the remaining DEGs had an absolute log fold change greater than 1 (Fig. 2A, volcano plot). In the endometrium, we detected the expression of 12043 and 12637 genes in both areas (caruncular (CAR) and inter-caruncular (ICAR)) on gestation days 18 and 25, respectively. In addition, we inferred 1963 and 2101 genes with differential transcript abundance between day 18 and day 25 CAR and ICAR areas, respectively (FDR<0.01, Figs. 2B, 2C), and 1178 DEGs were common between both areas of the endometrium (Dataset S2-S3).

**Fig. 2.**
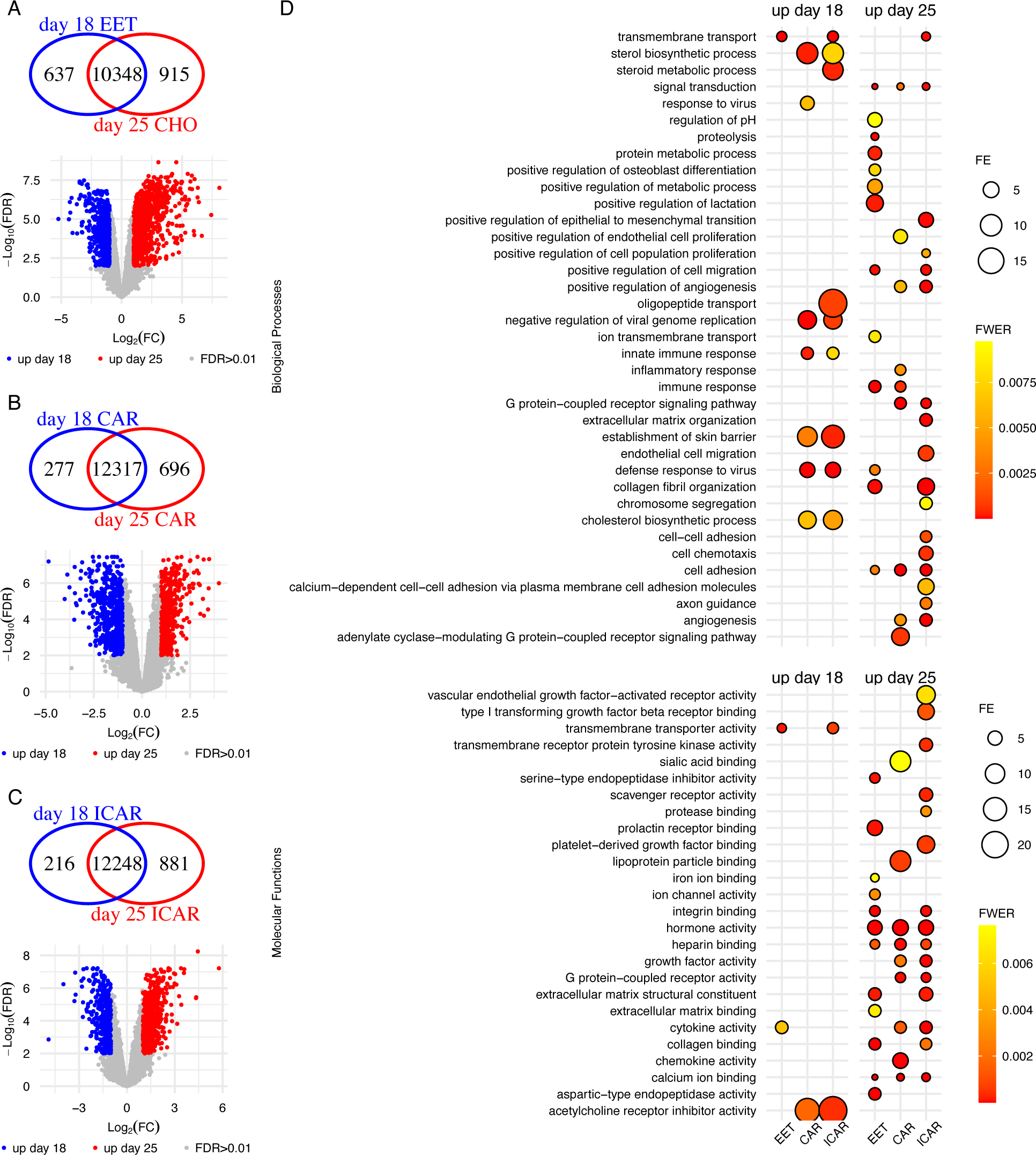
Developmental changes in the transcriptome of pregnancies derived by artificial insemination. (A) Extraembryonic tissues, (B) Caruncular, and (C) Inter-caruncular areas of the endometrium. (D) Summary of the gene ontology enrichment of the DEGs. Within each panel, the veen diagram indicates genes exclusively expressed in either group. The volcano plots show genes present in both groups tested, with genes showing quantitative differential transcript abundance shown in either red or blue. Only categories with FWER<0.01 are plotted on panel D. Graphs were produced from Datasets S1-S15. EET: extraembryonic tissues; CHO: chorion; CAR: caruncular; ICAR: inter caruncular; FE: fold enrichment; FWER: family-wise error rate.

Gene ontology enrichment analysis of those genes with greater transcript abundance on day 18 EET (N=2032) highlighted the biological process “transmembrane transport” as significantly enriched (FWER=9.7×10^-11^, N genes=65, Fig 2D, Dataset S4). The molecular function categories “transmembrane transporter activity” (N genes=27), “cytokine activity” (N genes=12) were significantly enriched (FWER<0.01), and “signaling receptor binding” (N genes=28) showed a tendency for significance (FWER<0.02) among 2032 genes with greater expression day 18 EET (Fig 2D, Dataset S5). Gene ontology enrichment analysis of genes with greater transcript abundance in day 25 chorion (N genes=1344) highlighted several biological processes as significantly enriched (FWER<0.01, Fig 2D, Dataset S6). Some examples of prominent biological importance are: “immune response” (N genes=28), “proteolysis” (N genes=86), “signal transduction” (N genes=109), “positive regulation of cell migration” (N genes=46), “defense response to virus” (N genes=28), and “cell adhesion” (N genes=48). Molecular categories also emerged as significantly enriched (FWER<0.01, Fig 2D, Dataset S7), for example, “hormone activity” (N genes=18), “integrin binding” (N genes=29), “serine-type endopeptidase inhibitor activity” (N genes=23), and “extracellular matrix structural constituent” (N genes=15). Although not significantly enriched among the DEGs, several other genes were annotated to biologically relevant gene ontology categories. For example, there were 30 and 54 DEGs annotated with the function of “DNA-binding transcription factor activity” with greater abundance on day 18 EET and day 25 chorion, respectively (Datasets S5, and S7).

Gene ontology enrichment analysis of genes with greater transcript abundance on day 18 CAR (N=805) or ICAR (N=583) regions of the endometrium showed an overlap of highly significant (FWER<0.01, Fig 2D, Datasets S8, and S9) categories: “defense response to virus”, “negative regulation of viral genome replication”, “sterol biosynthetic process”, “innate immune response”, “establishment of skin barrier”, and “cholesterol biosynthetic process”. Enrichment of the category “transmembrane transport” was highly significant in ICAR and had a strong tendency in CAR areas of the endometrium (FWER<0.01, Fig 2D, Datasets S8, and S9). The molecular function “acetylcholine receptor inhibitor activity” was significantly enriched in both areas of the endometrium (FWER<0.01, Datasets S10, and S11), whereas “transmembrane transporter activity” was significantly enriched in ICAR areas of the endometrium, and “monooxygenase activity” was significantly enriched (FWER=0.01) in CAR areas of the endometrium (Datasets S10, and S11).

The enrichment analysis of genes with greater transcript abundance on day 25 CAR (N=1158) or ICAR (N=1518) regions of the endometrium showed less overlapping of highly significant (FWER<0.01, Fig 2D, Datasets S12, and S13) categories: “angiogenesis”, “cell adhesion”, “G protein-coupled receptor signaling pathway”, “positive regulation of angiogenesis”, and “signal transduction”. Notably the categories “immune response”, “inflammatory response” and “positive regulation of endothelial cell proliferation” were enriched in CAR regions of the endometrium. By comparison, some of the biologically relevant categories that were enriched in ICAR regions of the endometrium were: “cell chemotaxis”, “cell-cell adhesion”, “endothelial cell migration”, “extracellular matrix organization”, “positive regulation of epithelial to mesenchymal transition”, and “transmembrane transport” (FWER<0.01, Fig 2D). Highly significant molecular functions that emerged in both CAR and ICAR areas of the endometrium were: “calcium ion binding”, “cytokine activity”, “G protein-coupled receptor activity”, “growth factor activity”, “heparin binding”, “hormone activity”, and “integrin binding” (FWER<0.01, Fig 2D, Datasets S14, and S15).

### Transcriptome alterations in pregnancies derived from *in vitro-produced* embryos in the implantation window relative to gestations initiated by artificial insemination

Next, we sought to determine the differences in transcript abundance associated with whether the embryo was produced *in vitro* or derived *in vivo*. We identified 885 genes with significant differences in transcript abundance in day 18 EETs (FDR<0.01, Fig. 3A, Dataset S16). The biological processes “cell communication”, “extracellular matrix organization”, “immune response”, and “proximal/distal pattern formation” were significantly enriched among the 721 genes with greater transcript abundance in conceptus originated by AI (FWER<0.01, Dataset S17). There was no enrichment among the genes with greater transcript abundance in conceptuses originating from the transfer of an *in vitro* produced embryo.

**Fig. 3.**
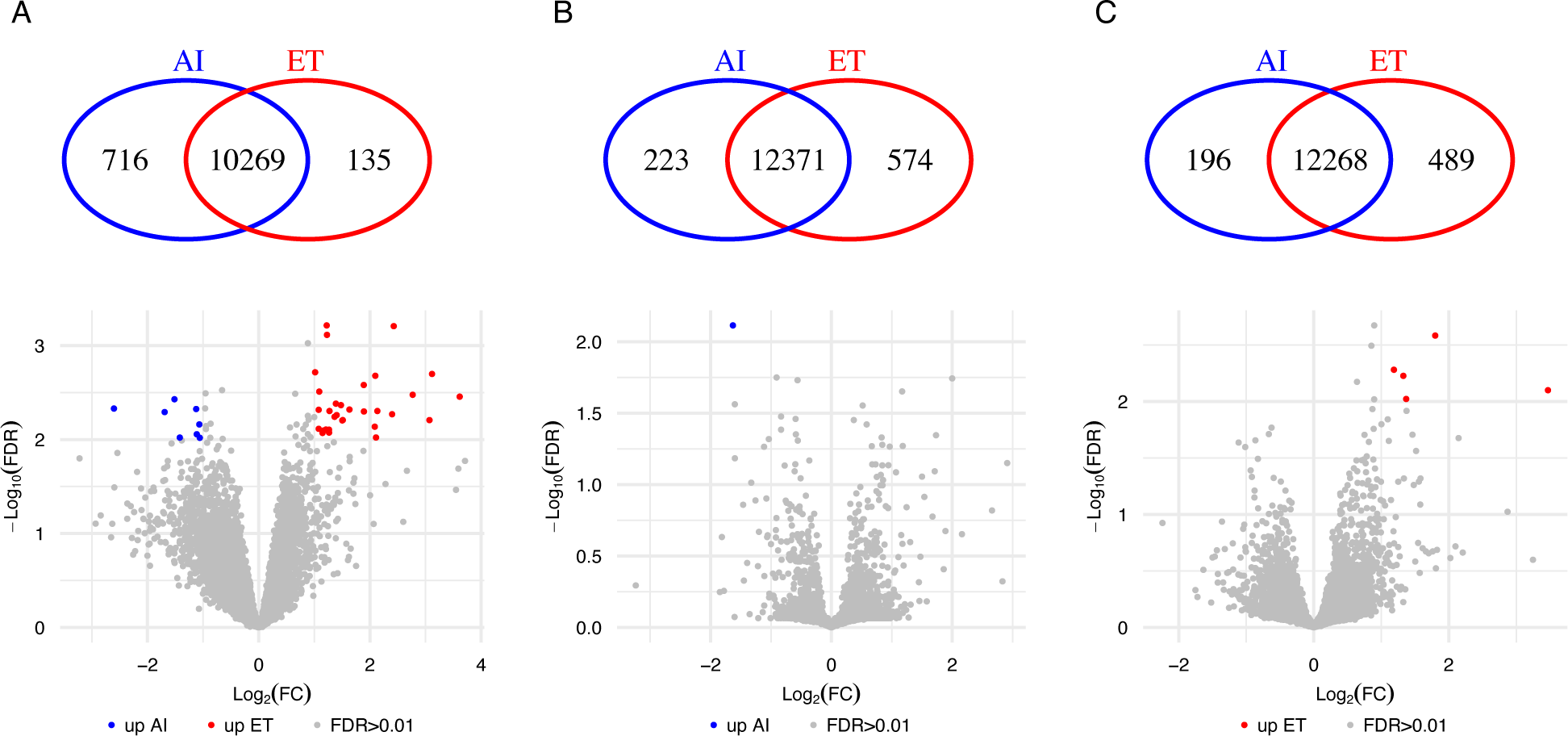
Differential transcript abundance related to *in vitro* versus *in vivo* produced embryos on gestation day 18. (A) EET, (B) Caruncular, and (C) Inter-caruncular areas of the endometrium. Within each panel, the veen diagram indicates genes exclusively expressed in either group. The volcano plots show genes present in both groups tested, with genes showing quantitative differential transcript abundance shown in either red or blue. AI: artificial insemination; ET: embryo transfer; FC: fold change; FDR: false discovery rate.

We determined 798 and 689 genes with significant differences in transcript abundance in CAR and ICAR areas of the endometrium, respectively, in relationship to the origin of the conceptus (FDR<0.01, Figs. 3B and 3C, Datasets S18-S19), in the endometrium of pregnancies terminated on gestation day 18. The biological process “ion transport” was significantly enriched (N genes= 9, FWER=0.0014) among the 196 genes with greater transcript abundance in ICAR areas of the endometrium harboring an AI-derived embryo (Datasets S20). On the other hand, the biological process “cilium movement” was significantly enriched (N genes=6, FWER=0.008) among the 493 genes with greater transcript abundance in ICAR areas of the endometrium harboring an *in vitro* produced embryo (Datasets S21). It was also notable that there were several genes annotated as “regulation of transcription by RNA polymerase II” in both CAR (N genes=45, FWER=0.02) and ICAR (N genes=36, FWER=0.12) areas of the endometrium harboring an *in vitro* produced embryo.

We identified 669 DEGs when comparing the transcriptome of chorion samples collected from *in vivo* or *in vitro* induced pregnancies terminated on gestation day 25 (FDR<0.01, Fig 4A, Dataset S22). We determined 725 and 728 genes with significant differences in transcript abundance in CAR and ICAR areas of the endometrium, respectively, in relationship to the origin of the conceptus (FDR<0.01, Figs. 4B and 4C, Datasets S23-S24) in the endometrium of pregnancies terminated on gestation day 25.

**Fig 4.**
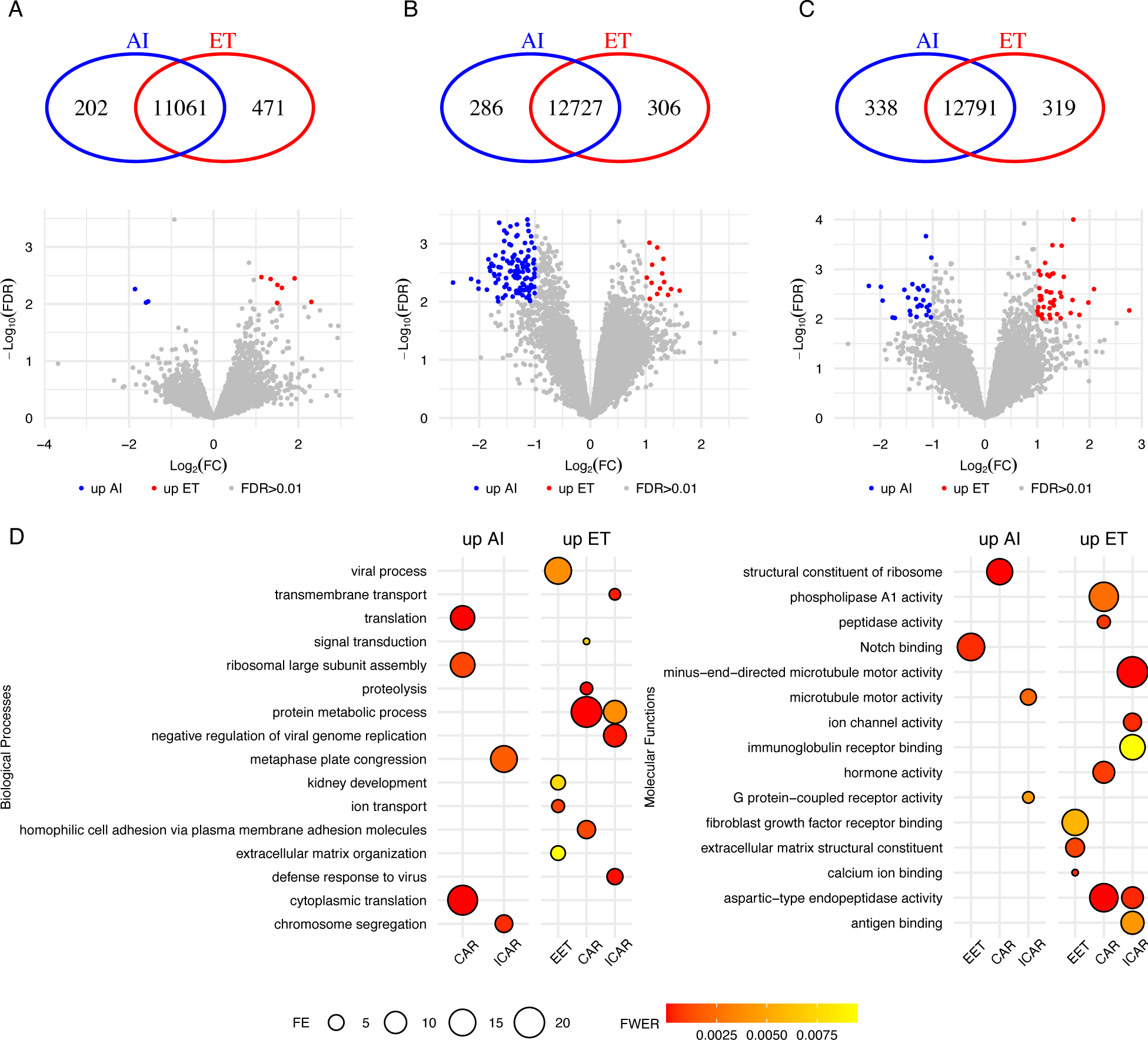
Differential transcript abundance related to *in vitro* versus *in vivo* produced embryos on gestation day 25. (A) Chorion, (B) Caruncular, and (C) Inter-caruncular areas of the endometrium. (D) Summary of the gene ontology enrichment of the DEGs. Within each panel, the veen diagram indicates genes exclusively expressed in either group. Thevolcano plots show genes present in both groups tested, with genes showing quantitative differential transcript abundance shown in either red or blue. Only categories with FWER<0.01 are plotted on panel D.

Among the 464 genes with greater transcript abundance in chorion from conceptuses derived *in vitr*o, we identified the following biological processes significantly enriched (FWER<0.01, Fig. 4D): “extracellular matrix organization”, “ion transport”, and “viral process”. Other biologically relevant categories such as “immune response”, “positive regulation of angiogenesis”, “transmembrane transport”, and “vascular endothelial growth factor receptor signaling pathway” showed a tendency for significance (FWER<0.05, Dataset S25). The molecular functions significantly enriched among these 464 genes were: “calcium ion binding”, “extracellular matrix structural constituent”, and “fibroblast growth factor receptor binding” (FWER<0.01, Fig. 4D, Dataset S26). By comparison, “Notch binding” was the molecular function enriched among the 205 genes with greater abundance in chorion from AI-derived conceptuses (FWER<0.01, Fig. 4D, Dataset S27).

Among the genes with greater abundance in endometrium harboring a day 25 conceptus produced *in vitro*, we identified “protein metabolic process” as the biological process common to both CAR (320 genes) and ICAR (363 genes) areas (FWER<0.01, Fig. 4D, Datasets S28, S29). The categories “homophilic cell adhesion via plasma membrane adhesion molecules”, “proteolysis”, and “signal transduction” (FWER<0.01, Fig. 4D) are biological processes significantly enriched in CAR areas of the endometrium. Interestingly, “defense response to virus”, and “negative regulation of viral genome replication” are biological processes also significantly enriched in ICAR areas of the endometrium. Several molecular functions were also significantly enriched among these genes (FWER<0.01, Fig. 4D, Datasets S30, S31).

Among the genes with greater abundance in endometrium harboring a day 25 conceptus produced by AI, most of the DEGs in CAR (129 out of 405 genes) were annotated to the biological process “translation” (FWER<0.01, Fig. 4D, Dataset S32). By contrast, in ICAR there only a few genes were annotated to the biological processes “chromosome segregation” and “metaphase plate congression” (FWER<0.01, Fig. 4D, Dataset S33). Similarly, a few molecular functions were also significantly enriched among these genes (FWER<0.01, Fig. 4D, Datasets S34, S35).

### Disturbed gene regulatory interactions between conceptus and endometrium in pregnancies derived from *in vitro* produced embryos relative to gestations initiated by artificial insemination

Given that we analyzed both the conceptus and the endometrium, we interrogated the data for genes expressing ligands and receptors with previously recorded interactions. On gestation day 18, we identified 223 ligands in the conceptus’ EET with the potential to form 796 interactions with 305 receptors in across both CAR and ICAR regions of the endometrium, 17 and five receptors in CAR or ICAR, respectively. We identified 279 ligands across both CAR and ICAR regions of the endometrium, 27 and three ligands in CAR or ICAR, respectively, with potential interactions with 267 receptors in the conceptus’ EET (SI Appendix Figs S1-S2, Datasets S36, S37). Related to how the conceptus was derived, in pregnancies originating from the transfer of an *in vitro* produced embryo, we did not detect transcripts for the genes coding for the ligands APOB, COL1A2, PLAT, PLG and the genes coding for the receptors CD44, IL17RB, IL17RB, ITGA2, NRP1 in EET. We also did not detect transcripts for the genes coding for the ligands CNTN2, COL11A1, IL17B, SPP1 and the genes coding for receptors CD36, ITGAM in the endometrium.

On gestation day 25 we identified 259 ligands in the chorion with the potential to form 897 interactions with 329 receptors in the endometrium (CAR and ICAR regions) and 12 receptors in CAR or ICAR regions of the endometrium. We also detected 302 ligands in the endometrium (CAR and ICAR regions), 21 and eight in CAR or ICAR, respectively, with the potential to interact with 283 receptors in the chorion (SI Appendix Figs S3-S4, Datasets S38, S39). Interestingly, no genes coding for ligands or receptors were differentially expressed due to *in vitro* embryo production.

Next, we tested for co-expression between the conceptus and endometrium with emphasis on genes that were exclusively detected in pregnancies harboring conceptuses derived *in vivo* or *in vitro*. We focused on the pairs of genes with correlated transcript abundances lower than -0.95 or greater than 0.95 (P≤0.001, Datasets S40, S41). In pregnancies initiated by AI and terminated on day 18, there were 59 genes expressed in the EET with strong co-expression with 46 genes in CAR (Fig 5A), and 45 genes expressed in the EET with strong co-expression with 41 genes in ICAR areas of the endometrium (Fig 5B). Notably, the transcripts of these genes were not detected in pregnancies initiated by the transfer of an *in vitro* produced embryo. Beyond that, we detected several, mostly positive, co-expressing pairs of genes that were formed in pregnancies initiated by the transfer of an *in vitro* produced embryo (14 genes in EET and 26 genes in CAR (Fig 5C); 32 genes in EET and 72 genes in ICAR (Fig 5D)).

**Fig. 5.**
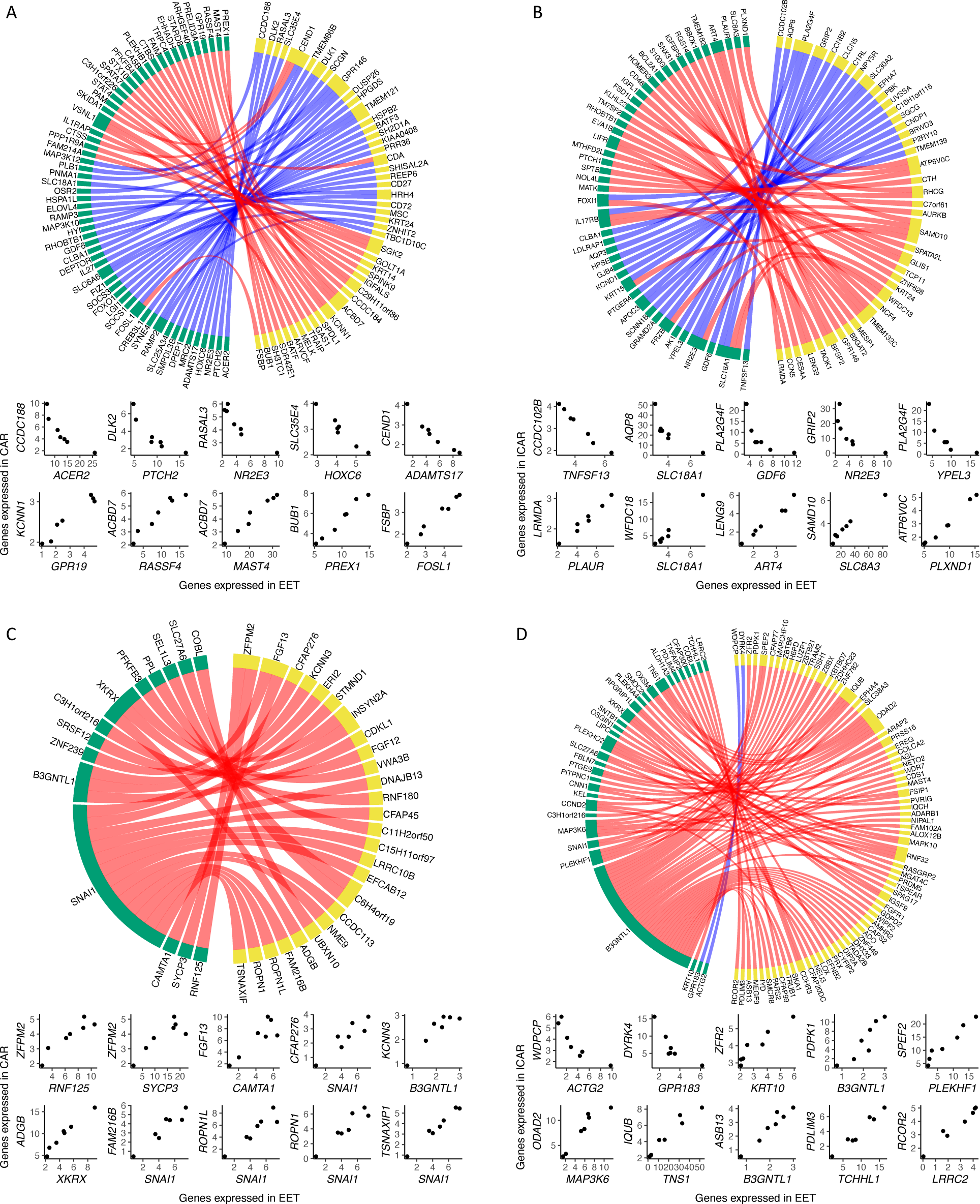
Co-expression networks present between conceptus and endometrium on gestation day 18. EET and (A) CAR or (B) ICAR from pregnancies initiated by artificial insemination. EET and (C) CAR or (D) ICAR from pregnancies initiated by the transfer of an *in vitro* produced embryo. In each panel, genes expressed in EET are represented in yellow and genes expressed in endometrium are represented in green. At the bottom of each network, there are scatterplots of representative gene pairs to illustrate their quantitative correlation. Only genes annotated with a symbol are depicted on these graphs.

We also evaluated the co-expression between conceptus and endometrium on gestation day 25 (|r|>0.95, P≤0.001, Datasets S42, S43). In pregnancies initiated by AI, there were 46 genes expressed in the chorion with strong co-expression with 34 genes in CAR (Fig 6A), and 57 genes expressed in the chorion with strong co-expression with 48 genes in ICAR (Fig 6B) areas of the endometrium. All co-expressing gene pairs did not have counterparts in the samples obtained by pregnancies initiated by the transfer of an *in vitro* produced embryo. In parallel, there were 129 genes expressed in chorion forming co-expressing pairs with 112 genes in CAR (Fig 6C) and 84 genes expressed in chorion forming co-expressing pairs with 78 genes in ICAR (Fig 6D) in pregnancies interrupted gestation day 25 after the transfer of an *in vitro* produced embryo.

**Fig. 6.**
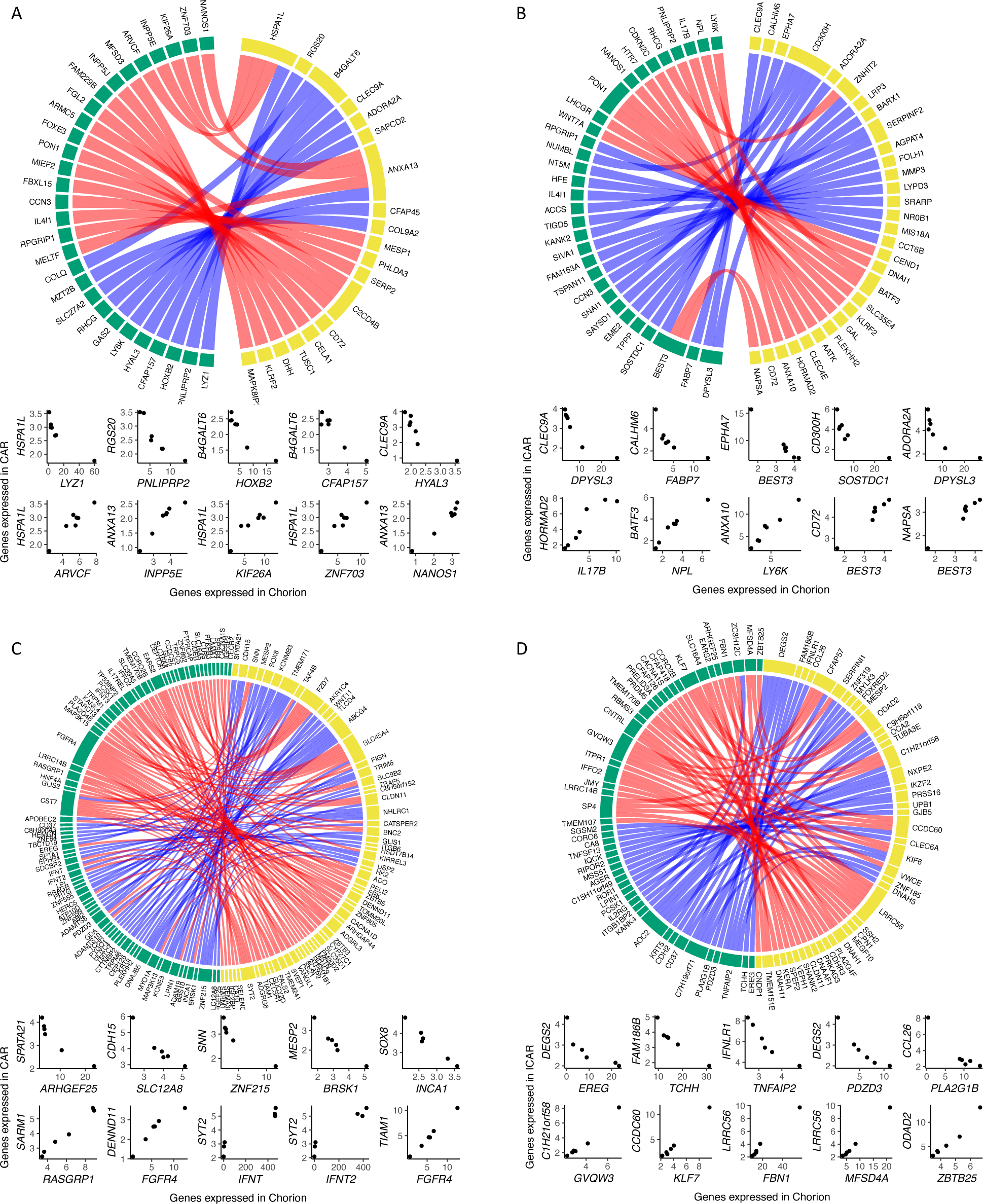
Co-expression networks present between conceptus and endometrium on gestation day 25. Chorion and (A) CAR or (B) ICAR from pregnancies initiated by artificial insemination. Chorion and (C) CAR or (D) ICAR from pregnancies initiated by the transfer of an *in vitro* produced embryo. In each panel, genes expressed in chorion are represented in yellow and genes expressed in endometrium are represented in green. At the bottom of each network, there are scatterplots of representative gene pairs to illustrate their quantitative correlation. Only genes annotated with a symbol are depicted on these graphs.

## DISCUSSION

Our study aimed to investigate the effects of *in vitro* embryo production systems on the molecular interaction between conceptus and endometrium during the attachment window. To achieve this goal, we conducted a comprehensive analysis of paired conceptus and endometrial tissues at two specific time points: the beginning of attachment (11) and the onset of epitheliochorial placentation (10). Our results provide strong evidence that *in vitro* production of cattle embryos has a profound impact on gene regulation in both the conceptus and endometrium. Moreover, this dysregulation is exacerbated by a loss of gene regulatory connections and the formation of new interactions between the conceptus and endometrium. Our findings have important implications for understanding the missing links and new interactions between the conceptus and endometrium during the establishment of a pregnancy resulting from artificial reproductive technologies, which may have adverse effects on gestation health.

Our work has some limitations. First, the model used compared pregnancies initiated by artificial insemination versus those initiated by the transfer of an in vitro-produced embryo, which might add layers of complexity to the comparisons. For instance, an experiment initiated by the transfer of in vivo derived embryos versus the transfer of in vitro produced embryos would have eliminated potential differences caused by allogenic conceptuses, or potential effects of the endometrium undergoing pharmacological estrous synchronization. Another limitation is that our data does not allow us to determine which cells produce the specific RNAs that emerged in our results. Further studies confirming the presence of the proteins for specific RNAs are also warranted to strengthen the findings. The findings, however, provide a systemic view of the alterations in the transcript abundance in conceptus and endometrium caused by in vitro production of embryos.

### Attachment window

We focused on a window of attachment that encompasses the beginning of the adhesion of the trophoblast to the epithelium of the endometrium (41). The trophoblast establishes a firm adhesion with the epithelium in much of its extension, including with the formation of papillae that invade the glandular lumen (41). By day 25, there is extensive interdigitation of microvilli between the trophoblast cells and the endometrial epithelia (11, 42). The dramatic change in the regulation of gene expression in both the EET and the endometrium between gestation days 18 and 25 reflects this dynamic and progressive interaction between the extraembryonic tissue and endometrium.

Among those genes with greater transcript abundance in the EET on day 18 of gestation initiated by artificial insemination versus day 25, there was an enrichment of genes from the solute carrier family, along with other genes involved in the transport of micro-molecules. This enrichment is well aligned with the need for trophoblast to utilize the amino acids (43, 44), glucose (45), and other nutrients for protein synthesis and cell proliferation (46). Notably, our results also showed a high number of genes annotated as having “cytokine activity” and/or “signaling receptor binding”. These results support the idea that Interferon tau c1 (*IFN-tau-c1*), which is part of the interferon-tau genes highly expressed between gestation days 17 and 19 (47), members of the Wnt family (48, 49), and other genes compose a series of genes driving complex signaling from the extraembryonic tissue to the endometrium.

In the endometrium of day 18 gestations initiated by artificial insemination relative to day 25, we observed several genes whose expression reflects the presence of the conceptus, including several that are inducible by the Interferon gene family (CAR: *IFI44L*, *IFI6*, *IFIH1*, *IFIT2*, *IFIT3*, *IFIT5*, *IRF6*, *IRF7*, *IRF9*, *ISG15*, *ISG20*; ICAR: *IFI44L*, *IFI6*, *IFIH1*, *IFIT3*, *IFNLR1*, *IRF6*, *IRF7*, *IRF9*, *ISG15*, *ISG20*). It was notable that the presence of the conceptus triggers genes directed to two major biological processes. First, an immune-related response (“defense response to virus”, “innate immune response”) is triggered across the endometrium. The genes identified in our analysis are most likely associated with an environment more permissive to T helper type 2 cells (50-52). Second, there were 31 genes associated with “transmembrane transport” significantly enriched in ICAR areas of the endometrium, and 32 genes tending towards significance in CAR (FWER=0.0159) areas of the endometrium. Although the glands are responsible for producing histotrophs (46, 53) and they are only present in the ICAR areas of the endometrium, the whole endometrium seems to have a role in the transport of nutrients to the lumen. During the period of conceptus initiation of attachment, our analysis highlighted genes that have critical roles in the functional remodeling of the endometrium (54) as part of uterine increased receptivity to the conceptus.

The genes with greater transcript abundance in chorion from conceptus produced in vivo on gestation day 25, relative to day 18, provide critical biological insights into the drivers of critical biological processes involved in the onset of placentation. For instance, “immune response” was the most significant biological process, composed of 28 genes. Interestingly, we identified the expression of *BOLA-A* and *BOLA-DMA*, which are potential mediators of immune tolerance (55) and the immunosuppressor Interleukin 1 receptor antagonist (56). It is also notable that the genes *CCL26* (57) and *ENPP2* (58) are involved in the modulation of immune functions, while also contributing to the migration and adhesion of the chorion to the endometrium. These genes are probably of critical importance for the modulation of the immune response in the endometrium, although these genes likely act indirectly, probably through the maintenance of an endometrial environment that is more favorable for anti-inflammatory T helper 2 cells (59, 60). While avoiding an attack from the immune system, several genes involved in proteolysis are likely to have a role in the remodeling of the endometrium which includes the removal of anti-adhesive glycoproteins (61) and also the removal of epithelial cells (11). Our analysis identified several other genes that are of great importance for the tight adhesion and signaling between the chorion and endometrium. Collectively we identified a catalog of genes with greater expression in the chorion on gestation day 25 with critical functions in the modulation of the onset of placentation.

An intensive cellular interaction between the conceptus and the endometrium is expected on gestation day 25 (11, 42) relative to earlier gestation days. Indeed, “cell adhesion” was the top significant biological process in both the CAR (47 genes) and ICAR (62 genes) areas of the endometrium. On this gestation day, the endometrium remains highly responsive to the presence of the conceptus because several genes also emerged in the highly significant biological process “signal transduction”. The biological process “immune response” was significant in the CAR areas of the endometrium, although with a different profile of genes relative to those observed on gestation day 18. On the other hand, many other biological processes were significantly enriched in the ICAR areas of the endometrium, such as “extracellular matrix organization”, “positive regulation of cell migration”, “positive regulation of cell population proliferation”, “positive regulation of cell migration”, “transmembrane transport”, and “cell chemotaxis”. All these categories show the contribution of the glandular area of the endometrium for the modulation of key functions that drive cellular proliferation and differentiation of the EET.

### Disrupted gene regulation in pregnancies derived by an *in vitro* production system

A remarkable finding from our transcriptome analysis was the detection, or lack thereof, of transcripts for several genes in samples obtained from pregnancies originated by the transfer of an *in vitro*-produced embryo. These results are a strong indication of a massive alteration in gene regulation caused by *in vitro* culture of embryos.

The number of genes and DEGs (ET vs AI) obtained for extraembryonic tissue collected on gestation day 18 is in alignment with a previous study conducted with embryos produced by somatic cell nuclear transfer and in vitro culture (22). An interesting finding was that the extraembryonic tissue from *in vitro-produced* conceptuses had more genes with loss of expression rather than a gain of expression. One potential reason for this result is a dysregulation in the methylation profiles of promoters caused by *in vitro* culture (62) that may lead to a genome-wide increase in methylation, thus transcription repression. Another possibility is that few drivers of gene transcription (i.e.: 26 genes associated with “negative regulation of transcription by RNA polymerase II” or 41 genes associated with “positive regulation of transcription by RNA polymerase II”) have repressed transcription at the chromatin level and precipitate a cascade of transcription repression. The extraembryonic tissue of conceptuses produced *in vitro* and collected on gestation day 18 lacked transcripts from several genes involved in biological processes essential for attachment (i.e.: “extracellular matrix organization”, “cell communication”, “cell communication”). The category “signal transduction”, with 40 genes, is also of extreme importance for communication with the endometrium, although this category did not reach our significance threshold. The enrichment of genes in these categories, along with others of critical importance for intra-uterine development, is likely a strong indication of the mechanisms that lead to critical losses during early pregnancy.

The enrichment of “cilium movement”, among the genes with greater expression in ICAR areas of the endometrium harboring a day-18 *in vitro* produced conceptus, was intriguing. A recent report indicated a reduction of ciliated cells on the surface of ICAR areas with the progression of the estrous cycle (day 14 vs day 0, (63)). Although we did not evaluate ciliated cells on the surface of the endometrium, one could speculate that these genes (*CFAP221*, *DNAH11*, *DNAH5*, *ODAD2*, *ODAD3*, *TTC29*, *ZBBX*) are upregulated in ICAR because of a dysregulated interaction with the *in vitro* produced conceptus. This is an example that the remodeling of the endometrium can also be altered at the cellular level, and not only at the molecular level. At the molecular level, we observed a series of ion transport genes downregulated in ICAR areas, which is likely to lead to a reduced transfer of nutrients from the uterine glands to the lumen. Also at the molecular level, although not significantly enriched (FWER=0.026 in CAR, FWER=0.12 in ICAR), there were 31 and 22 annotated genes associated with “regulation of transcription by RNA polymerase II” upregulated in CAR and ICAR, respectively, areas of the endometrium harboring *in vitro* produced conceptuses. This might be associated with a cascade of dysregulated gene expression, presented elsewhere (64), as an adverse ripple of the altered interaction of the conceptus and endometrium (21) that has consequences throughout pregnancy.

An intriguing result emerged from our enrichment analysis of biological processes of genes upregulated in chorion from day 25 conceptuses produced *in vitro* versus day 25 conceptuses generated by artificial insemination. There were seven genes associated with “ion transport” (*SLC12A6*, *SLC12A8*, *SLC30A10*, *SLC4A7*, *TRPC5*, *TRPM1*, *TRPM6*) and four genes associated with “viral process” (ENSBTAG00000048696, ENSBTAG00000050000, ENSBTAG00000051140, ENSBTAG00000054418) that were also upregulated in the extraembryonic tissue on gestation day 18, when compared to chorion collected on day 25 of conceptuses produced by artificial insemination. One possible explanation is that the chorion of day 25 conceptuses produced *in vitro* is morphologically delayed relative to those produced *in vivo*. However, we did not process conceptuses with morphological signs of delayed development nor signs of tissue degeneration for transcriptome analysis. Thus, all the conceptuses that we used for chorion sampling and transcriptome analysis presented similar developmental stages, and their chorion tissues were morphologically similar when observed under the stereoscope. An alternative explanation is that these conceptuses produced *in vitro* have dysregulated gene transcription (22), and thus certain groups of genes would remain transcriptionally active. Along with the possibility of dysregulated gene expression, we observed the down-regulation of four “Notch binding” genes (*CCN3*, *CHAC1*, *DLL4*, *JAG2*) that are extremely important for the downstream regulation of genes in trophoblast cells, with implications in development and differentiation (65).

Also supporting the idea of dysfunctional dysregulation of gene activity, we observed that ICAR areas of the endometrium harboring a day 25 conceptus produced *in vitro* had eight annotated genes (*DDX58*, *IFI44L*, *IFIH1*, *IFNLR1*, *MX1*, *RSAD2*, *ZBP1*, *ZNFX1*) associated with “defense response to virus” and seven annotated genes (*AQP8*, *CACNA1S*, *CLCN5*, *OCA2*, *SLC45A2*, *SLC9B1*, *SLCO4C1*) associated with “transmembrane transport”. Both categories were also enriched in ICAR areas on gestation day 18 versus day 25 when harboring conceptuses produced by artificial insemination. This result is an indication that a series of genes do not reduce their transcriptional activity in ICAR areas when the conceptus is produced *in vitro*.

### Molecular interaction between conceptus and endometrium in pregnancies derived by an *in vitro* production system

Previous studies demonstrated that the EET and the endometrium interact at the molecular level in two dimensions. In one dimension, there are ligands and receptors (21, 22), and in another dimension, there are gene regulatory networks forming between the conceptus and the endometrium (23). Our study allowed us to analyze both dimensions of this intricate interaction.

Our comprehensive transcriptome analysis identified a large catalog of genes coding for ligands and receptors with transcripts detected in the EET, chorion, and endometrium. It was surprising that a limited number of genes coding for ligands and receptors were differentially expressed due to the origin of the conceptus. Notably, two of the ligands lacking in the EET from conceptus produced *in vitro*, plasminogen (*PLG*) and plasminogen activator (*PLAT*), may have an important impact on the interaction with the endometrium. Bovine pre-implantation embryos produce plasminogen and plasminogen activator even after hatching (66), and the abundances of their transcripts are not negligible in EET derived by artificial insemination (4.1 and 47.3, respectively). Transcripts of plasminogen were also detected in the trophectoderm of day 15.75 pig conceptuses (67). Although the context of interaction between trophoblast and endometrium has not been investigated in cattle, plasminogen and the activation system have critical roles in cell signaling, extracellular matrix degradation, and the migration of cells from the immune system (reviewed in (68)). In mice, plasminogen and the activation system participate in endometrial remodeling and trophoblast migration into the uterine wall (67). The lack of these two proteins in the EET of day 18 conceptuses produced *in vitro* may result in impaired adhesion to the endometrium.

Our quantitative analysis of genes expressed in both compartments revealed an extensive rearrangement of gene co-expression between the conceptus and the endometrium. On both gestation days 18 and 25, this rearrangement involved the loss and creation of several interactions due to the presence of an *in vitro* produced conceptus. One of the most notable observations was the co-expression between *IFNT*, *IFNT2* and *IFNT3* expressed in chorion with genes expressed in the CAR areas of the endometrium (IFNT:*NHLRC1*, *IFNT*:*SYT2*, *IFNT2*:*NHLRC1*, *IFNT2*:*SYT2*, *IFNT3*:*ARHGAP44*, *IFNT3*:*SYT2*). This interaction was mostly caused by three chorion samples that still expressed two genes of the interferon tau family, and their expression this late in the development is also in line with the molecular delay we previously discussed. The quantitative interaction of genes expressed in the conceptus and endometrium is a phenomenon we have identified previously (23), but our current findings indicated that this molecular connection is severely impacted by the origin of the conceptus and reinforces the idea that the endometrium adapts to dramatic changes in gene expression caused by artificial reproductive technologies (22, 38, 40).

In closing, our work has identified several alterations in gene expression caused by *in vitro* production of embryos during the peri-attachment period. One of the features associated with implantation failure during this window of pregnancy is the delayed trophoblast elongation (29). Our results contributed evidence that the chorion of day 25 conceptuses produced *in vitro* has a delay in the regulation of gene expression. Many of the genes with transcripts in the chorion of day 25 conceptuses produced *in vitro* were present in the EET of day 18 conceptuses produced by artificial insemination. It is highly likely that the EET of day 18 conceptuses produced *in vitro* have a delayed regulation of gene expression, but our experimental design did not permit us to test this hypothesis. The alterations in gene expression in conceptuses caused by *in vitro* production are reflected in the endometrium which also responds to the stimuli (22, 38, 40), probably in a delayed manner. Collectively, this dysfunctional interaction between some of the conceptuses produced *in vitro* and the endometrium may have a role in the interruption of a proportion of pregnancies.

## MATERIAL AND METHODS

### *In vitro* production of embryos and cryopreservation

All chemicals were obtained from Sigma-Aldrich (St. Louis, MO) or Fisher (Pittsburgh, PA), unless otherwise stated. Embryos were produced following established protocols and media described in detail elsewhere (69-71). Cumulus-oocyte complexes were aspirated from follicles (3-8mm in diameter) in abattoir-derived ovaries. We washed the cumulus-oocyte complexes in Tissue Culture Medium-199 with Hanks salts supplemented with 25mM HEPES followed by *in vitro* maturation in Tissue Culture Medium-199 with Earle salts (Gibco, Grand Island, NY) supplemented with 10% fetal bovine serum, 100 IU/ml penicillin, 100 μg/ml streptomycin, 0.2 mM sodium pyruvate, 2 mM L-glutamine, 50 ng/ml recombinant human epidermal growth factor (Invitrogen, Waltham, MA), and 5 μg/ml of follicle-stimulating hormone (Bioniche Animal Health, Athens, GA). In vitro maturation was carried out for 22-24 hours at 38.5°C in a humidified atmosphere containing 5% CO_2_.

*In vitro* matured cumulus-oocyte complexes were washed trice with HEPES-TALP and placed in IVF-TALP for *in vitro* fertilization. Sperm from a single sire was prepared by centrifugation in a gradient of Isolate (Irvine Scientific) and washed twice by centrifugation in SP-TALP. Sperm was added to the fertilization dish at the concentration of 1×10^6^/ml, followed by the addition of penicillamine-hypotaurine-epinephrine solution. *In vitro* fertilization was carried out for 17-19 hours at 38.5°C in a humidified atmosphere containing 5% CO_2_. Cumulus cells were removed from putative zygotes by vortexing in 400 μl/of HEPES-TALP. Putative zygotes were then cultured in groups of up to 50 in 500 μl of SOF-BE2, covered with 300 μl of light mineral oil. *In vitro* culture was carried out at 38.5°C in a humidified atmosphere containing 5% CO_2_ and 5% O_2_. Seven days after *in vitro* fertilization, grade 1 (72) blastocysts were cryopreserved using the slow-freezing procedure in ethylene-glycol solution (73).

### Estrous synchronization, artificial insemination, and embryo transfer

We used nulliparous heifers (15-19 months of age, weighing >296 kg) of Angus-cross genetic background for this experiment. Animals were randomized into one of the four experimental groups, based on whether they would be artificially inseminated or serve as recipients for embryo transfer, and based on whether the pregnancy would be terminated on gestation days 18 or 25.

We started estrous synchronization (74) by inserting a controlled internal drug release (CIDR, 1.38 g progesterone), which was removed after 14 days. Sixteen days post removal of CIDR we administered prostaglandin F 2 alpha (25mg, Lutalyse, Zoetis) and animals were observed for estrus. If the heifer was assigned to be a recipient, we administered gonadotropin-releasing hormone (100 mcg, Cystorelin, Cystorelin, Merial, Athens, GA) 66 hours post-prostaglandin administration. The heifers assigned to be artificially inseminated were inseminated 12-16 hours after onset of estrus or a few exceptions were inseminated at 66 hours post-prostaglandin administration and administered gonadotropin-releasing hormone (100 mcg, Cystorelin, Cystorelin, Merial, Athens, GA). All heifers were inseminated to one sire, which was the same sire used for *in vitro* embryo production.

For embryo transfer, seven days post-administration of gonadotropin-releasing hormone, we evaluated the presence of a corpus luteum by ultrasonography. If a corpus luteum was present, one embryo was deposited in the uterine horn ipsilateral to the corpus luteum. If the heifer did not present a corpus luteum, she was re-enrolled in estrous synchronization.

### Sample collection

Heifers were euthanized with captive bolt on day 18 or 25 of pregnancy. The reproductive tract was removed from each heifer within 15 minutes of euthanasia and immediately prepared for flushing. To prepare the reproductive tract for flushing, first, we removed the mesometrium from the uterine horns. Next, we cut the base of the uterine body to remove the cervix. Then, we inserted an 18g needle coupled to a syringe into the cranial portion of the ipsilateral horn and flushed 20 ml of nuclease-free phosphate-buffered saline solution towards the base of the uterine body.

On gestation day 18, conceptuses were collected on a cell strainer, immediately transferred to a solution of RNALater (ThermoFisher, Vilnius, Lithuania), and maintained on ice until sectioned for long-term freezing at -80°C. On gestation day 25, conceptuses were transferred from the phosphate-buffered saline solution into RNALater and maintained on ice until dissection. On the same day of collection, with the aid of a stereoscope, we dissected the EET and snap-frozen samples in liquid nitrogen followed by long-term freezing at -80°C

Following the flushing, we opened the ipsilateral horn with a longitudinal section and rinsed the endometrium with approximately 50 ml of nuclease-free phosphate-buffered saline solution. Using fine dissection scissors, we sampled caruncular and intercaruncular areas of the endometrium, within 2 mm of the lumen to prevent sampling of the deep stroma or myometrium. We collected samples of caruncular and intercaruncular areas across the uterine horn. At the collection site, samples were immediately placed in tubes and snap-frozen in liquid nitrogen followed by long-term freezing at -80°C.

### RNA extraction and production of sequencing data

The sections selected for RNA extraction were separated, frozen in liquid nitrogen, and ground with mortar and pestle. We mixed the tissue with 800 μl of TRIzol Reagent (75) (Ambion, Carlsbad, CA), and proceeded with a standard guanidinium thiocyanate-phenol-chloroform extraction (76, 77) following the manufacturer’s protocol. Sequencing libraries were prepared at the Genomics Technology Core at the University of Missouri using the TruSeq Stranded mRNA kit (Illumina, San Diego, CA). Approximately 35 million pair-end sequences of 100 nucleotides long were produced per sample in a NovaSeq6000 instrument (Illumina, San Diego, CA) at the Genomics Technology Core at the University of Missouri.

### Alignment, processing of sequences, and gene quantification

We aligned the reads to the bovine genome (Bos_taurus.ARS-UCD1.2.104) using Hisat2 (v 2.2.1) (78). Next, we used Samtools (v 1.10) (79) to retain the reads with one match to the genome, followed by the removal of duplicates using bammarkduplicates from biobambam2 (v 2.0.95) (80).

We used featurecounts (v 2.0.1) (81) to count reads according to the ensemble annotation (Bos_taurus.ARS-UCD1.2.104) (82, 83). We quantified counts per million (CPM) and fragments per kilobase per million (FPKM), using the functions “cpm” or “rpkm” from the “edgeR” package (84, 85). We also calculated transcript per million using the formula presented elsewhere (86). We retained genes annotated as protein-coding, pseudogenes, or long non-coding RNA, and only used genes for further analysis if CPM and FPKM were >1 in at least five samples within each group. This approach was adopted to reduce the number of genes with highly variable expression and thus prone to producing confounding results (87).

### Alignment, processing of sequences, and transcript abundance for sexing

We used a pipeline published elsewhere (88) to align sequences produced from EET to the sequence of cattle chromosome Y (GenBank: CM011803.1) (89). Briefly, the fasta sequence of chromosome Y was appended to the cattle genome fasta sequence followed by the creation of an index and alignment of reads using Hisat2 (v 2.2.1) (78). We also used featurecounts to count the reads on chromosome Y using the annotation of chromosome Y provided by Liu and colleagues (89). Samples that had reads assigned to the genes “ENSBIXG00000029788”,“ENSBIXG00000029774” (*ZRSR2*) and “ENSBIXG00000029763” (*OFD1*) were inferred as males (88).

### Analysis of differential transcript abundance

We compared transcript abundance between samples from each group using the R packages ‘edgeR’ (84, 85), with the quasi-likelihood test, and ‘DEseq2’ (90), using the Wald’s and likelihood test. For tests involving conceptuses, we added sex as a fixed effect in the model. The nominal P values of both tests were corrected for multiple hypothesis testing using the false discovery rate (FDR) method (91). Differential transcript abundance was assumed when FDR<0.01 for both tests and |Log fold change| > 1.

### Analysis of ligand-receptors

We obtained a comprehensive list of ligand-receptors published elsewhere (92). Only ligands and receptors that were annotated with the following gene ontology cellular compartment annotations were considered for further analysis: “plasma membrane”, “extracellular region”, “cell surface”, “cell-cell junction”, “extracellular space”, “extracellular matrix”, “cell-cell contact zone”, “plasma membrane region”, “plasma membrane protein complex”, “cell-substrate junction”, or “protein complex involved in cell-matrix adhesion”.

### Co-expression analysis

We adopted the procedures recommended in this benchmarking work by Johnson and Krishnan (93). For each group, first, we calculated the CPM metric using the trimmed mean of M values method (94) using the function “calcNormFactors” from the R package ‘edgeR’ (84, 85). Second, we transformed the CPM using the arcsine transformation 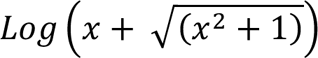 using the “asinh” function in R base. Next, we calculated the Pearson coefficient of correlation using the function “corAndPvalue” from the package ‘WGCNA’ (95) for each pair of genes with transcript abundance estimated in the extraembryonic tissue and endometrium.

### Enrichment of gene ontology

We carried out tests for enrichment or gene ontology categories using “goseq” package (96) in R software. In all tests, we used the genes whose transcript abundances were estimated for the samples being tested as the background. We adjusted the nominal P value for multiple hypothesis testing by controlling the familywise error rate following the method proposed by Holm (97) using the function “p.adjust” from the ‘stats’ R package. We inferred significance when FWER ≤ 0.01 and assumed a tendency for significance when 0.01 < FWER < 0.05.

## ACKNOWLEDGMENTS

We thank Mr. Barnes Wilborn and his team from The Lambert-Powell Meat Laboratory at Auburn University for their assistance in the collection of reproductive tracts.

## Data sharing plans

Raw data produced in the present research and metadata are deposited in the Gene Expression Omnibus repository under access: GSE232489. The code used for data analysis will be available from the authors upon request.

## Funding information

This project was supported by Agriculture and Food Research Initiative Competitive Grant no. 2018-67015-31936 from the USDA National Institute of Food and Agriculture.

## Notes

Competing Interest Statement: The authors declare no conflict of interest.

https://www.ncbi.nlm.nih.gov/geo/query/acc.cgi?acc=GSE232489

### Competing Interest Statement

The authors have declared no competing interest.

