## Appendix for "Extensive rewiring of the gene regulatory interactions between in vitro-produced conceptuses and endometrium during attachment"

**This PDF file includes:**

Table 1. Number of pairs of sequences produced for each sample analyzed.

Fig. 1. Connectome between ligands present in extra embryonic tissue and receptors present in endometrium of pregnancies initiated by artificial insemination and terminated on gestation day 18.

Fig. 2. Connectome between ligands present in endometrium and receptors present in extra-embryonic tissues of pregnancies initiated by artificial insemination and terminated on gestation day 18.

Fig. 3. Connectome between ligands present chorion and receptors present in endometrium of pregnancies initiated by artificial insemination and terminated on gestation day 25.

Fig. 4. Connectome between ligands present in endometrium and receptors present in extra-embryonic tissues of pregnancies initiated by artificial insemination and terminated on gestation day 25.

Fig. 5. Determination of the sex of the conceptuses using RNA-sequencing data mapped to genes on the cattle Y chromosome.

Supplementary text and associated figures

Fig. 6. Representative genes showing significant interaction in extraembryonic tissues between gestation day and source of conceptuses (in vitro produced embryos (ET) or artificial insemination (AI)).

Fig. 7. Representative genes showing significant interaction in caruncular areas of the endometrium between gestation day and source of conceptuses (in vitro produced embryos (ET) or artificial insemination (AI)).

Fig. 8. Representative genes showing significant interaction in inter-caruncular areas of the endometrium between gestation day and source of conceptuses (in vitro produced embryos (ET) or artificial insemination (AI)).

Table 1. Number of pairs of sequences produced for each sample analyzed.

| Heifer ID | Group day | Group source | Endometrium area | EET tissue | Reads produced | Reads assigned |
|---|---|---|---|---|---|---|
| 5 | D18 | AI | caruncular | - | 86,692,596 | 23,517,097 |
| 5 | D18 | AI | intercaruncular | - | 140,311,688 | 56,481,413 |
| 5 | D18 | AI | - | EET | 6,600,416 | 1,744,211 |
| 9 | D18 | AI | caruncular | - | 49,808,501 | 9,145,480 |
| 9 | D18 | AI | intercaruncular | - | 55,547,739 | 17,548,794 |
| 9 | D18 | AI | - | EET | 108,392,882 | 52,148,047 |
| 13 | D18 | AI | caruncular | - | 153,000,048 | 61,326,027 |
| 13 | D18 | AI | intercaruncular | - | 146,594,554 | 72,656,598 |
| 13 | D18 | AI | - | EET | 110,537,200 | 48,343,988 |
| 17 | D18 | AI | caruncular | - | 137,644,172 | 61,702,672 |
| 17 | D18 | AI | intercaruncular | - | 36,783,269 | 16,463,532 |
| 17 | D18 | AI | - | EET | 51,320,281 | 26,411,012 |
| 20 | D18 | AI | caruncular | - | 175,339,488 | 51,963,207 |
| 20 | D18 | AI | intercaruncular | - | 29,871,995 | 4,676,900 |
| 20 | D18 | AI | - | EET | 112,406,999 | 48,147,292 |
| 23 | D18 | AI | caruncular | - | 153,174,496 | 54,986,222 |
| 23 | D18 | AI | intercaruncular | - | 146,609,821 | 68,856,879 |
| 23 | D18 | AI | - | EET | 13,584,906 | 7,245,562 |
| 37 | D18 | AI | - | EET | 11,675,080 | 6,524,762 |
| 37 | D18 | AI | caruncular | - | 131,518,550 | 40,040,636 |
| 37 | D18 | AI | intercaruncular | - | 63,981,055 | 24,131,792 |
| 7 | D18 | ET | - | EET | 9,643,898 | 5,288,144 |
| 7 | D18 | ET | caruncular | - | 93,800,089 | 44,080,119 |
| 7 | D18 | ET | intercaruncular | - | 141,226,863 | 60,293,803 |
| 15 | D18 | ET | - | EET | 6,813,236 | 3,940,823 |
| 15 | D18 | ET | caruncular | - | 134,907,091 | 40,383,400 |
| 15 | D18 | ET | intercaruncular | - | 96,845,129 | 19,253,876 |
| 27 | D18 | ET | - | EET | 63,365,397 | 26,873,747 |
| 27 | D18 | ET | caruncular | - | 164,758,263 | 67,081,837 |
| 27 | D18 | ET | intercaruncular | - | 139,498,223 | 59,678,927 |
| 31 | D18 | ET | - | EET | 57,462,158 | 27,401,054 |
| 31 | D18 | ET | caruncular | - | 165,173,200 | 52,705,654 |
| 31 | D18 | ET | intercaruncular | - | 133,823,596 | 47,439,695 |
| 33 | D18 | ET | - | EET | 10,905,256 | 6,237,953 |
| 33 | D18 | ET | caruncular | - | 106,999,856 | 43,964,267 |
| 33 | D18 | ET | intercaruncular | - | 143,028,520 | 70,252,251 |
| 115 | D18 | ET | - | EET | 60,587,014 | 28,291,679 |

| | | | | | | |
|---|---|---|---|---|---|---|
| 115 | D18 | ET | caruncular | - | 102,475,229 | 51,740,673 |
| 115 | D18 | ET | intercaruncular | - | 147,735,842 | 74,489,271 |
| 169 | D18 | ET | - | EET | 44,731,366 | 22,690,915 |
| 169 | D18 | ET | caruncular | - | 152,929,028 | 72,667,262 |
| 169 | D18 | ET | intercaruncular | - | 142,833,759 | 70,719,022 |
| 1 | D25 | AI | - | chorion | 51,931,506 | 23,061,478 |
| 1 | D25 | AI | caruncular | - | 129,161,271 | 45,618,852 |
| 1 | D25 | AI | intercaruncular | - | 70,813,007 | 20,758,469 |
| 3 | D25 | AI | - | chorion | 69,366,087 | 33,119,544 |
| 3 | D25 | AI | caruncular | - | 111,335,812 | 46,519,614 |
| 3 | D25 | AI | intercaruncular | - | 27,122,671 | 13,376,952 |
| 18 | D25 | AI | - | chorion | 49,862,665 | 26,331,782 |
| 18 | D25 | AI | caruncular | | 139,755,892 | 40,364,497 |
| 18 | D25 | AI | intercaruncular | | 113,432,640 | 51,450,842 |
| 19 | D25 | AI | - | chorion | 113,745,258 | 50,985,167 |
| 19 | D25 | AI | caruncular | - | 91,317,951 | 30,415,957 |
| 19 | D25 | AI | intercaruncular | - | 56,525,758 | 23,612,236 |
| 39 | D25 | AI | - | chorion | 47,176,157 | 21,841,155 |
| 39 | D25 | AI | caruncular | - | 136,204,900 | 31,661,479 |
| 39 | D25 | AI | intercaruncular | - | 113,088,436 | 37,995,871 |
| 40 | D25 | AI | - | chorion | 57,272,489 | 28,885,204 |
| 40 | D25 | AI | caruncular | - | 126,703,771 | 40,861,469 |
| 40 | D25 | AI | intercaruncular | - | 128,076,942 | 54,830,852 |
| 44 | D25 | AI | - | chorion | 45,129,275 | 23,848,089 |
| 44 | D25 | AI | caruncular | - | 137,536,995 | 23,149,184 |
| 44 | D25 | AI | intercaruncular | - | 88,625,937 | 21,614,515 |
| 10 | D25 | ET | - | chorion | 99,981,806 | 20,836,766 |
| 10 | D25 | ET | caruncular | - | 149,246,368 | 57,453,057 |
| 10 | D25 | ET | intercaruncular | - | 123,941,731 | 52,918,838 |
| 28 | D25 | ET | - | chorion | 99,520,024 | 52,104,712 |
| 28 | D25 | ET | caruncular | - | 143,388,088 | 64,720,538 |
| 28 | D25 | ET | intercaruncular | - | 52,825,122 | 28,648,850 |
| 104 | D25 | ET | - | chorion | 41,477,030 | 21,388,322 |
| 104 | D25 | ET | caruncular | - | 156,675,060 | 64,754,573 |
| 104 | D25 | ET | intercaruncular | - | 134,694,483 | 68,600,021 |
| 107 | D25 | ET | - | chorion | 51,430,229 | 26,038,492 |
| 107 | D25 | ET | caruncular | - | 125,918,830 | 57,618,157 |
| 107 | D25 | ET | intercaruncular | - | 159,545,608 | 69,827,114 |
| 120 | D25 | ET | - | chorion | 43,417,109 | 22,852,471 |
| 120 | D25 | ET | caruncular | - | 158,567,116 | 74,295,707 |

| | | | | | | | |
|---|---|---|---|---|---|---|---|
| | 120 | D25 | ET | intercaruncular | - | 133,943,989 | 69,391,552 |
| | 162 | D25 | ET | - | chorion | 96,586,527 | 49,925,597 |
| | 162 | D25 | ET | caruncular | - | 289,255,911 | 128,807,044 |
| | 162 | D25 | ET | intercaruncular | - | 281,977,557 | 144,947,101 |
| Total | | | | | | 8,287,516,757 | 3,414,968,615 |
| Average | | | | | | 102,315,022 | 42,160,106 |

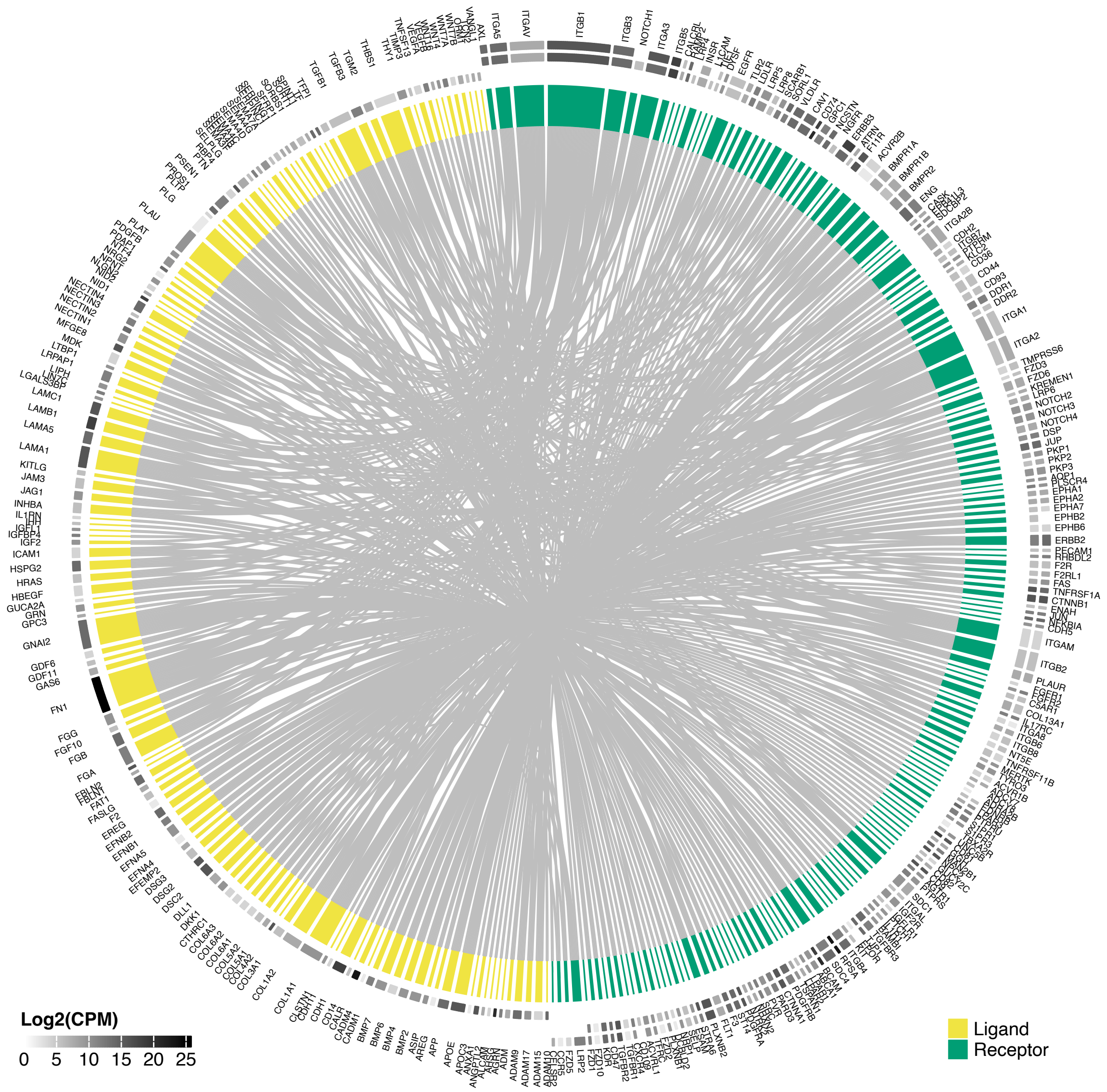


Fig. 3. Connectome between ligands present in extra embryonic tissue and receptors present in endometrium of pregnancies initiated by artificial insemination and terminated on gestation day 18. Inside and outside tracks of receptors represent transcript abundance in caruncular and inter-caruncular areas of the endometrium.

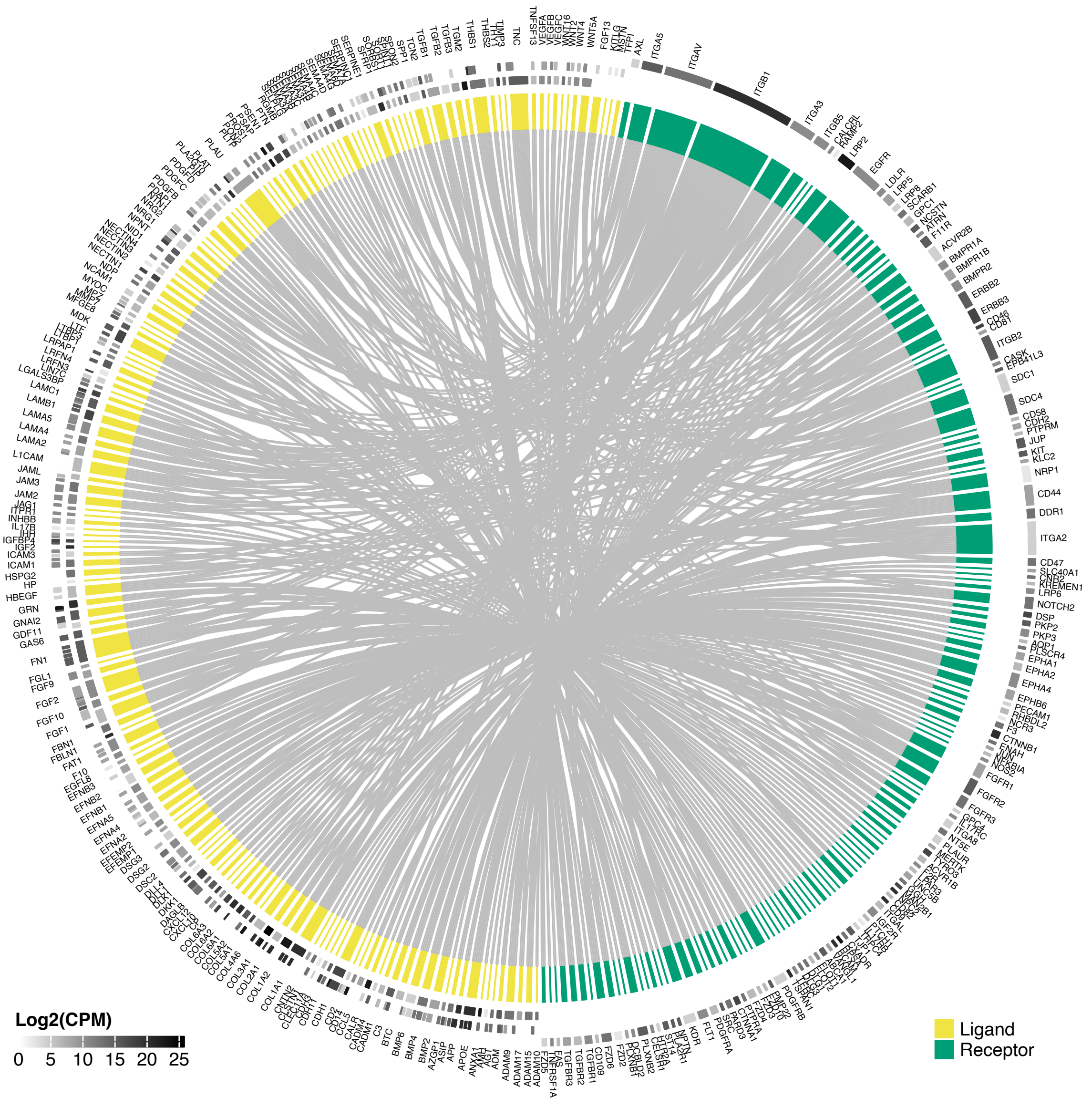


Fig. 4. Connectome between ligands present in endometrium and receptors present in extra-embryonic tissues of pregnancies initiated by artificial insemination and terminated on gestation day 18. Inside and outside tracks of ligands represent transcript abundance in caruncular and inter-caruncular areas of the endometrium.

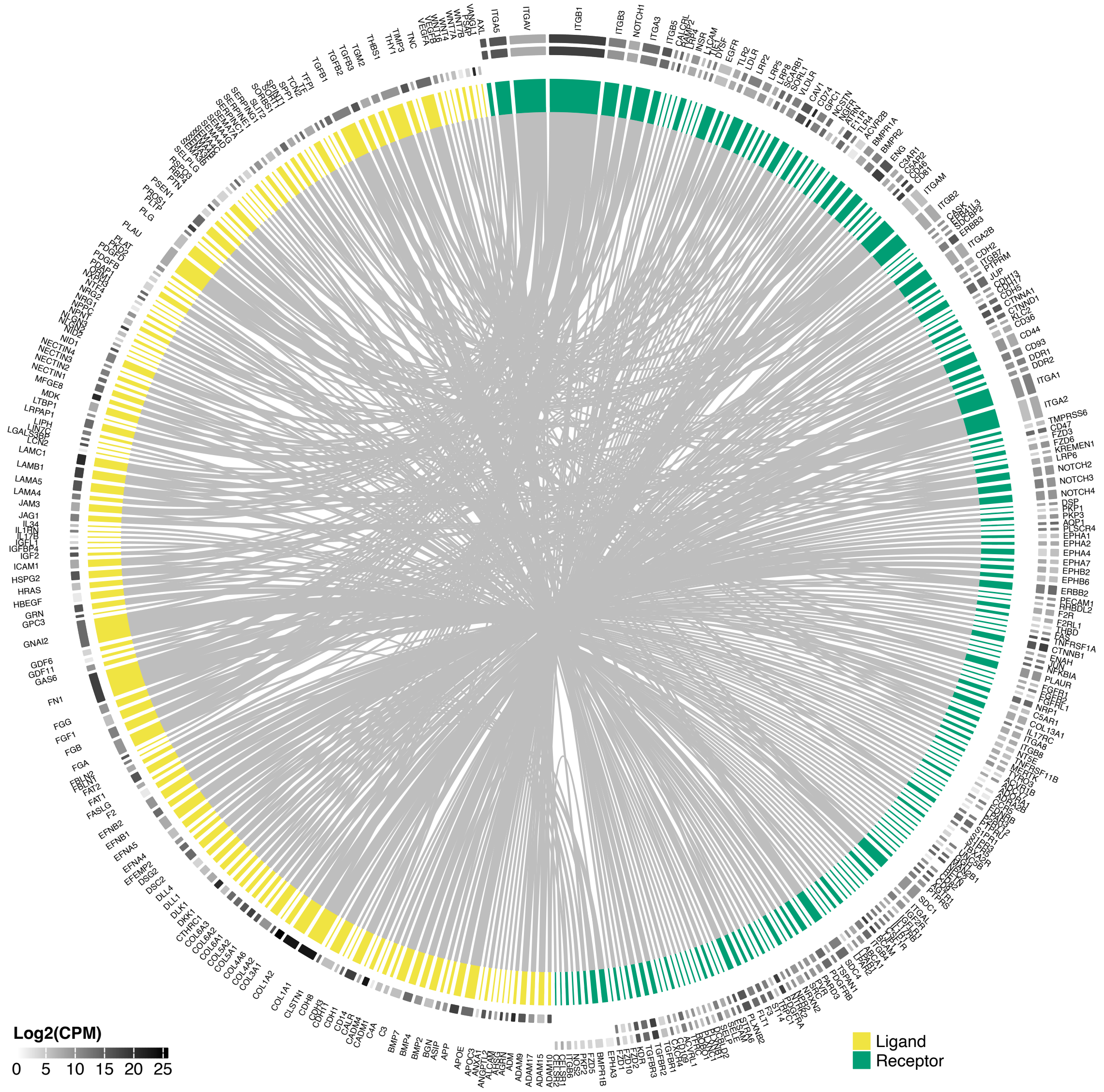


Fig. 3. Connectome between ligands present chorion and receptors present in endometrium of pregnancies initiated by artificial insemination and terminated on gestation day 25. Inside and outside tracks of receptors represent transcript abundance in caruncular and inter-caruncular areas of the endometrium.

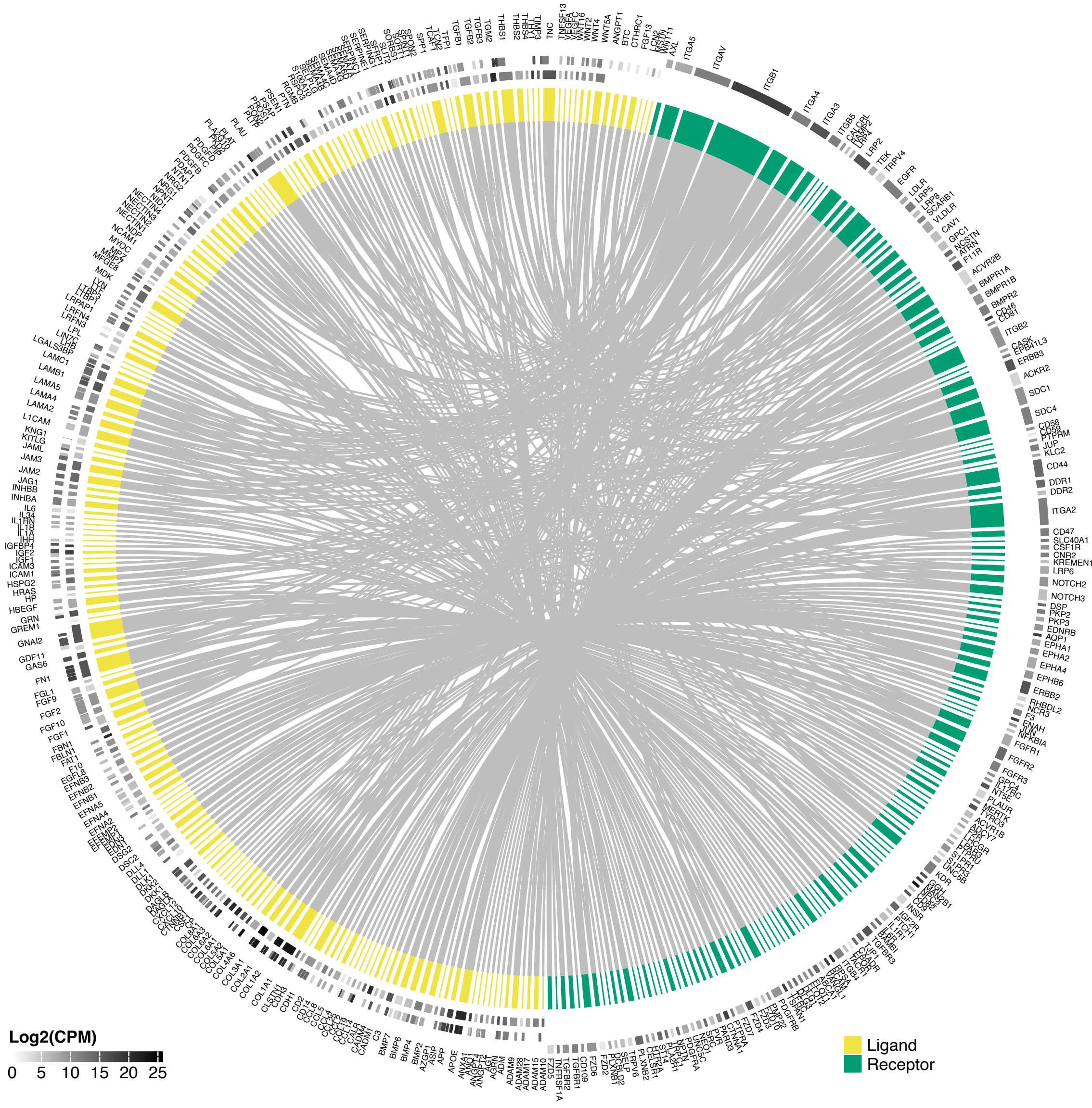


Fig. 4. Connectome between ligands present in endometrium and receptors present in extra-embryonic tissues of pregnancies initiated by artificial insemination and terminated on gestation day 25. Inside and outside tracks of ligands represent transcript abundance in caruncular and inter-caruncular areas of the endometrium.

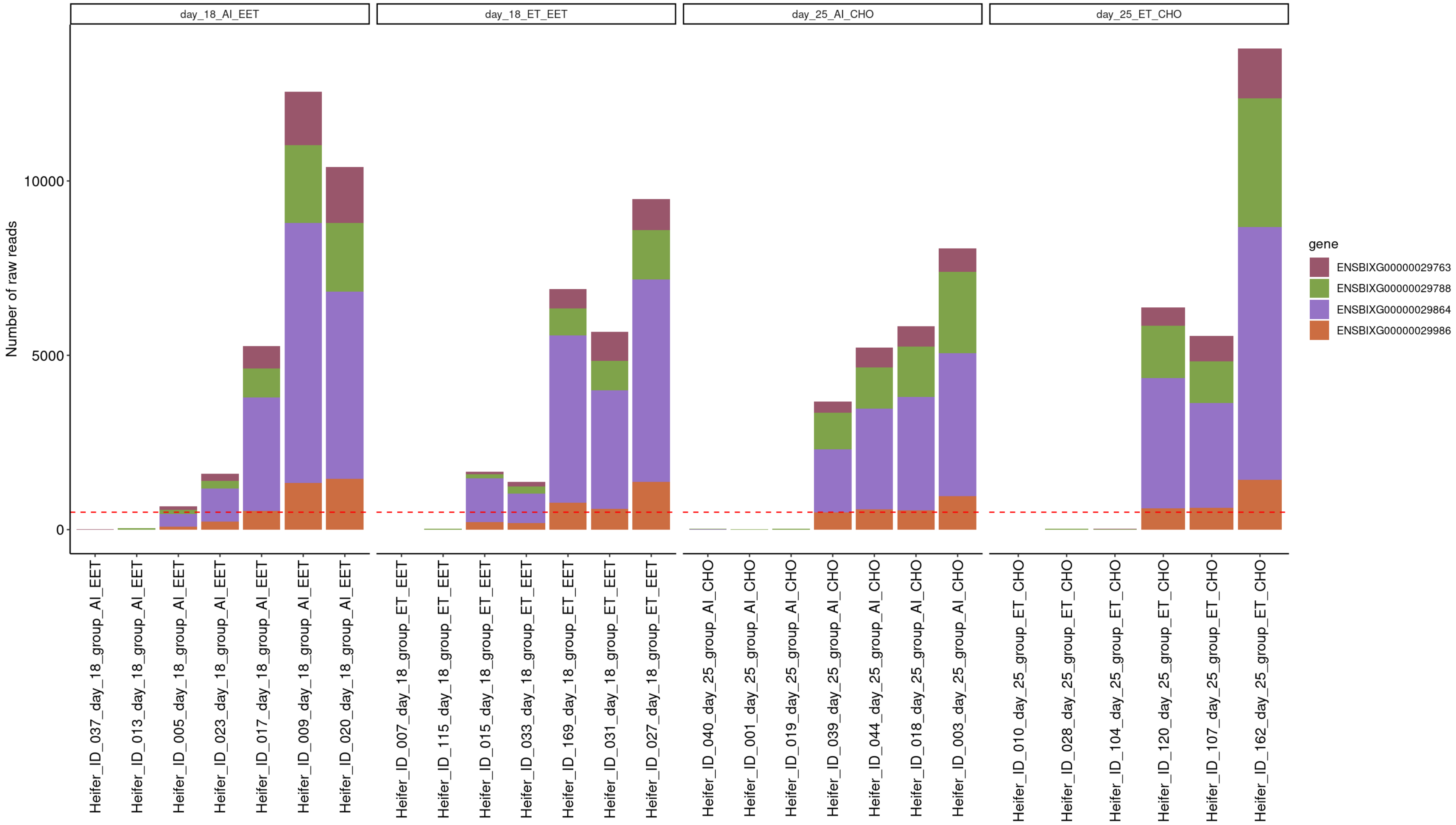


Fig. 5. Determination of the sex of the conceptuses using RNA-sequencing data mapped to genes on the cattle Y chromosome. Horizontal dashed line marks 500 reads mapped to four genes to determine sex. Samples with more than 500 reads assigned to these four genes were designated as male

### Supplementary text and associated figures

#### Changes in the transcriptome of extraembryonic tissues and endometrium between gestation days 18 and 25 in gestations initiated by the transfer of an *in vitro-produced* embryo

We inferred 3654 genes with differential transcript abundance between day 18 EET and day 25 chorion from conceptuses produced *in vitro* and transferred to a recipient on day seven (FDR<0.01, Dataset S44). There were 2407 and 648 genes with greater and lower transcript abundance in day 25 chorion compared to day 18 EET, respectively (FDR<0.01, Dataset S44). Analysis of the endometrial samples revealed 1361 and 1314 genes with differential transcript abundance between day 18 and day 25 caruncular and inter caruncular areas, respectively (FDR<0.01, Datasets S45 and S46).

We also analyzed the interaction between gestation days (18 and 25) and source of the conceptus (*in vivo* derived or *in vitro* produced). Notably, the analysis of extraembryonic tissues revealed 82 genes with significant interaction between gestation day and source of the conceptus and overlapped with differential transcript abundance between conceptuses of both sources on both gestation days (FDR<0.01, Dataset S47, see Fig S6 for example of 15 genes). The analysis of caruncular areas of the endometrium revealed 71 genes with significant interaction between gestation day and source of the conceptus and overlapped with differential transcript abundance between samples of endometrium harboring conceptuses of both sources on both gestation days (FDR<0.01, Dataset S48, see Fig S7 for example of 15 genes). The analysis of inter-caruncular areas of the endometrium revealed 53 genes with significant interaction between gestation day and source of the conceptus and overlapped with differential transcript abundance between samples of endometrium harboring conceptuses of both sources on both gestation days (FDR<0.01, Dataset S49, see Fig S8 for example of 15 genes).

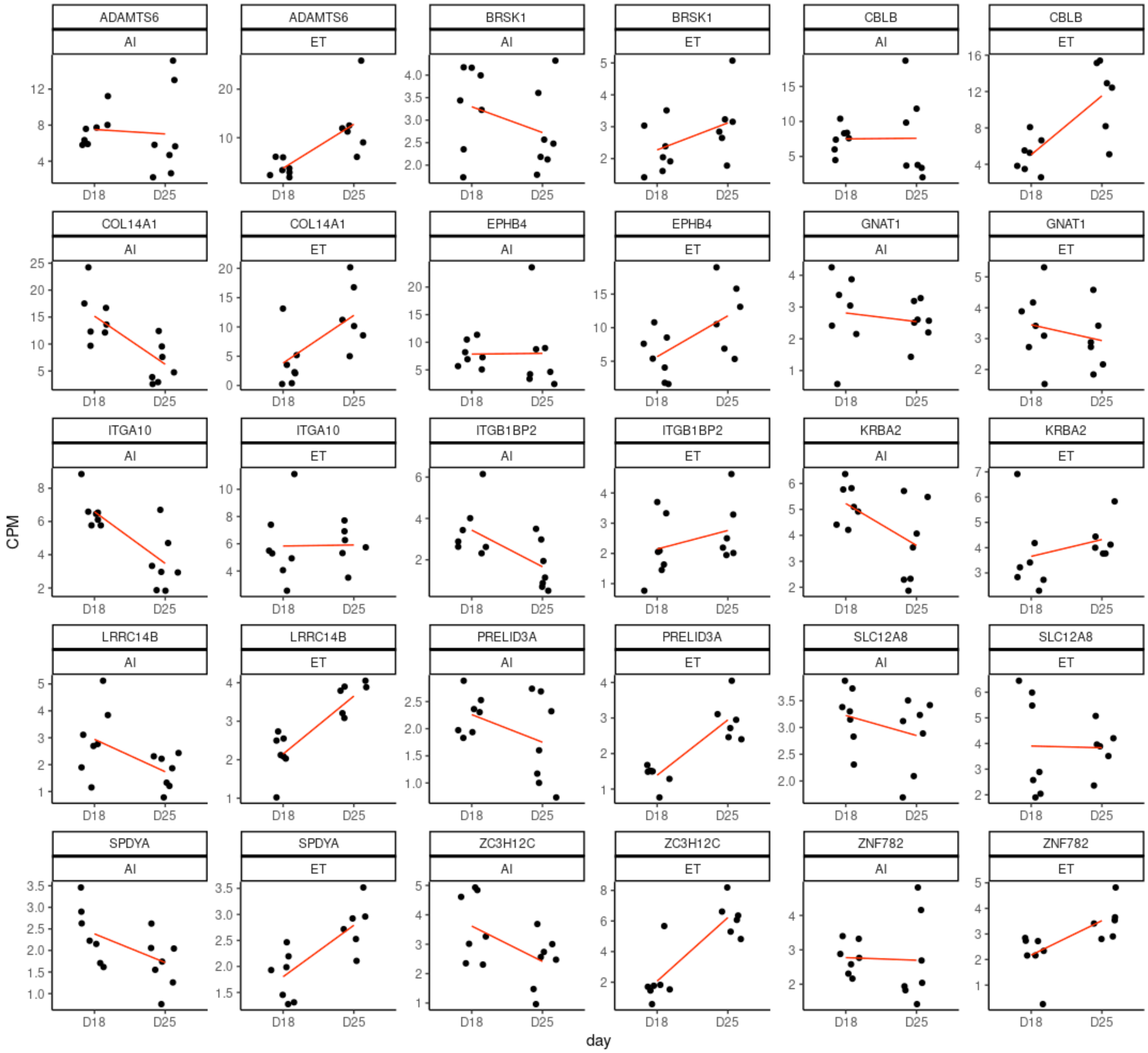


Fig. 6. Representative genes showing significant interaction in extraembryonic tissues between gestation day and source of conceptuses (in vitro produced embryos (ET) or artificial insemination (AI)).

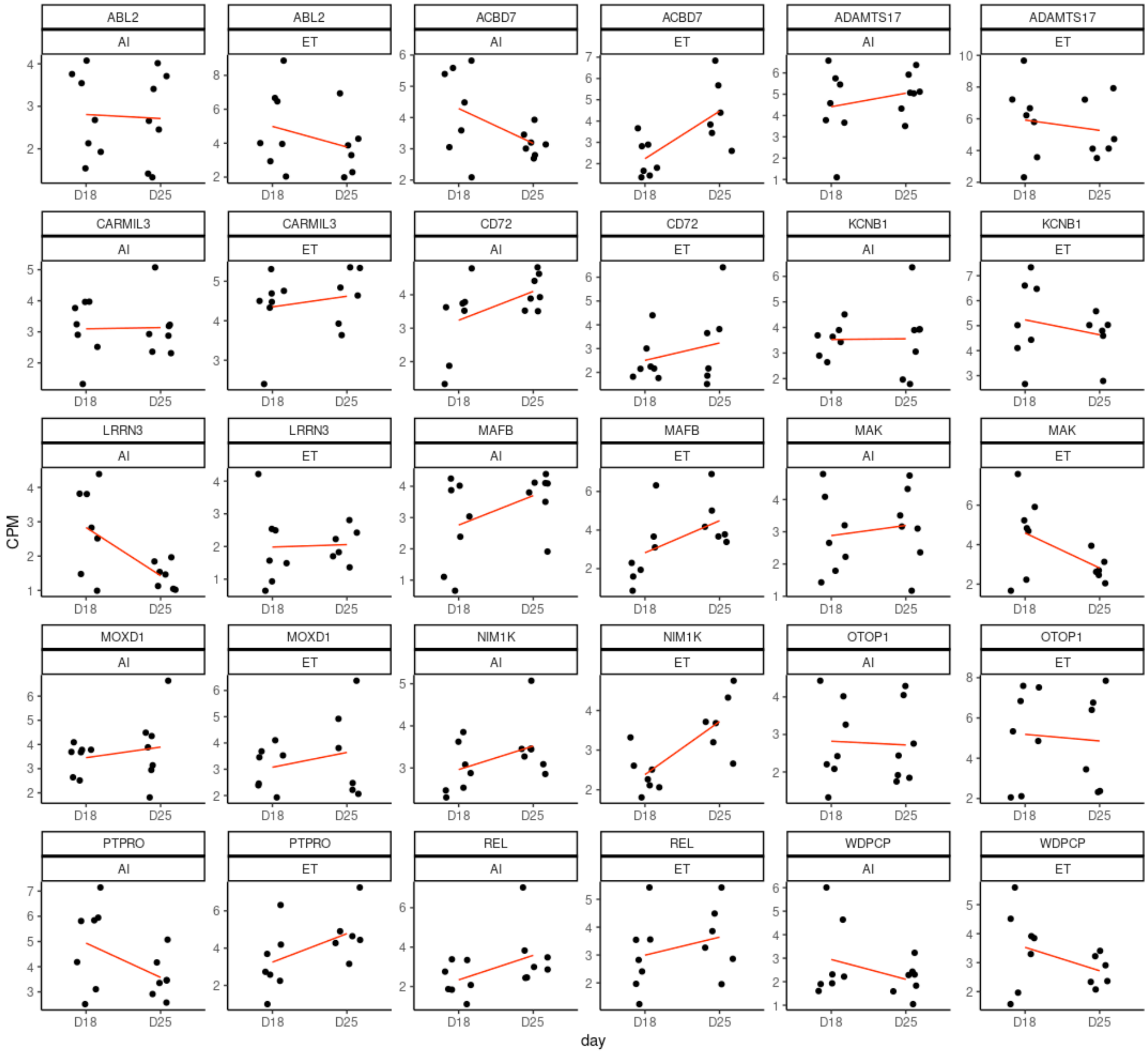


Fig. 7. Representative genes showing significant interaction in caruncular areas of the endometrium between gestation day and source of conceptuses (in vitro produced embryos (ET) or artificial insemination (AI)).

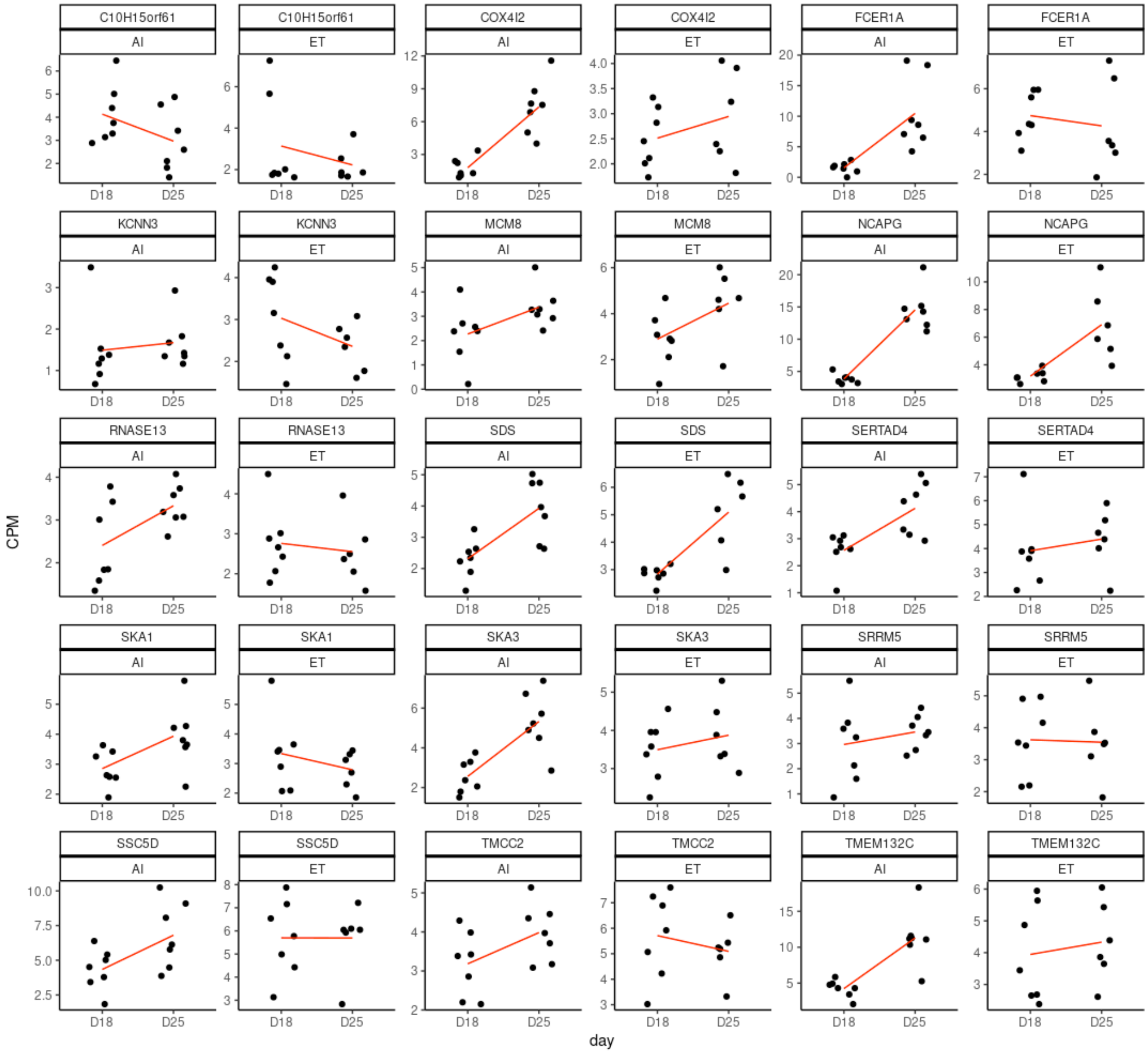


Fig. 8. Representative genes showing significant interaction in inter-caruncular areas of the endometrium between gestation day and source of conceptuses (in vitro produced embryos (ET) or artificial insemination (AI)).
